# Antibiotic-induced *Malassezia* spp. expansion in infants promotes early-life immune dysregulation and airway inflammation in mice

**DOI:** 10.1101/2024.04.24.590822

**Authors:** Erik van Tilburg Bernardes, Mackenzie W. Gutierrez, William N. T. Nguyen, Emily M. Mercer, Hena R. Ramay, Thaís Glatthardt, Carolyn A. Thomson, Tisha Halim, Nithya Gopalakrishnan, Kristen Kalbfleish, Kamala D. Patel, Kathy D. McCoy, Stephen B. Freedman, Marie-Claire Arrieta

## Abstract

Antibiotics have deleterious consequences for the gut microbiome and can increase the risk of childhood asthma. While the effects of antibiotics on the bacterial microbiome and asthma risk are well characterized, their impact on the fungal microbiome (mycobiome) remains vastly unexplored. We investigated the effect of antibiotic use on the gut mycobiome in an observational, prospective clinical study of young infants. Antibiotic treatment resulted in increased fungal abundance and expansion of the yeast *Malassezia* spp. Based on these findings, germ-free mouse pups were colonized with a defined consortium of mouse-derived bacteria (Oligo-MM12) with or without *Malassezia restricta*. Colonization with this yeast increased myeloid and lymphoid intestinal immune responses deemed critical in atopy development, and elevated airway inflammation in house-dust mite (HDM)-challenged mice. Further evaluation in eosinophil-deficient mice revealed that the observed immune response is partially dependent on this cell type. This translational work demonstrates that fungal overgrowth and expansion of *Malassezia* spp. are previously overlooked collateral effects of infant antibiotic use, which may offer a potential strategy to prevent or mitigate pediatric asthma and related conditions.

**One Sentence Summary:** Antibiotic-induced *Malassezia* spp. expansion in infants promotes early-life immune dysregulation and airway inflammation in gnotobiotic mice.

## MAIN

Perturbances to the microbiome early in life, known as dysbiosis, precede and are causally associated with immune dysregulation that can result in immune-mediated diseases, such as asthma^1–5^. Fungi constitute an integral part of the microbiome, and only recently have studies begun to investigate their role in influencing health outcomes in the host^6,7^. Population-based studies have identified altered mycobiome signatures associated with asthma risk ^8–12^. Some of these studies have further demonstrated causality in mouse models of airway inflammation ^8,9^. For instance, neonatal mice exposed to *Pichia kudriavzevii,* a fungus increased in infants at risk of asthma development ^9^, displayed increased airway inflammation following HDM challenge, with higher T helper type 2 (Th2) and Th17 cells infiltration in the lungs ^13^.

Fungal symbionts are critical for the induction of early tolerogenic immune responses in the host. This is not surprising considering that fungi are bonafide members of the microbiome, and evolutionary forces likely shaped the mammalian immune system to tolerate commensal fungal presence ^14^. For instance, commensal fungi induced migration of regulatory retinol dehydrogenase positive (RALDH^+^) dendritic cells (DC) to mesenteric and peripheral lymph nodes in germ-free mice ^15^, supporting gut-associated lymphoid tissue and mucosal tolerance development early in life. Fungal colonizers also promote the development of homeostatic effector T subsets both in the intestines and systemically, including Th17 cells, which promote barrier function and tune host immune responses ^7,16,17^. Yet, it remains unclear how perturbances to the gut mycobiome early in life can impact host immune development.

Antibiotic administration during early life is one of the most important dysbiosis-inducing factors, as it can profoundly disrupt the commensal microbiome ^18,19^. Antibiotics induce strong compositional and functional changes in both bacterial and fungal communities^18,19^, and are one of the strongest risk factors for allergic asthma development ^20–24^. In a case-control study in Ecuadorian infants, antibiotic use during the first year of life was associated with fungal overgrowth and increased susceptibility to allergic asthma ^9^. Antibiotic-induced mycobiome changes have also been linked to exacerbation of airway inflammation in animal models ^25–29^. While this may suggest that fungal overgrowth and/or dysbiosis may contribute to the antibiotic-induced microbiome alterations that cause immune dysregulation during infancy, the impact of early-life antibiotics on the mycobiome and its effects on host immune development remains unknown.

Here, we analyzed the compositional changes to the bacterial and fungal gut microbiome in a prospective clinical study of infants < 6 months of age receiving urgent antibiotic therapy. Upon finding that antibiotic use led to the expansion of *Malassezia* spp., we developed a gnotobiotic mouse model to study the causal contribution of a representative species from this genus in intestinal immune development and susceptibility to allergic airway inflammation. This comprehensive work provides novel understandings of the impact of antibiotics on infant microbiome composition and may offer a potential strategy to prevent or mitigate pediatric asthma and related conditions.

## RESULTS

### The ANTIBIO Study

The ANTIBIO study recruited 47 infants aged < 6 months evaluated in the emergency department of the Alberta Children’s Hospital (ACH-ED) who were prescribed parenteral (i.e. intravenous or intramuscular) antibiotics **(study design in Fig. 1A and *Supplementary Materials*)**. Study participants were separated into two groups based on treatment duration, short-term (receiving antibiotics for 2-3 days; *n=*22) and long-term (4-14 days; *n=*25). Antibiotic treatment duration included both parenteral antibiotics administered in the hospital and oral antibiotics prescribed at discharge (**Fig. 1a**). Study subject characteristics are presented in **Table S1**. Most variables were distributed similarly between the two treatment groups (*P >* 0.05), except for antibiotic treatment received, birth weight (median weight at birth: 3.4 kg for short-term vs. 3.0 kg for long-term; *P=*0.02), presence of pets in the household (no pets at home: 36.4 % for short-term vs. 68.0 % for long-term; *P=*0.03), and heart-rate at triage (mean heart-rate: 153 bpm for short-term vs. 175 bpm for long term group; *P=*0.007) **(Table S1)**. While the ANTIBIO study was not designed to examine the associations between antibiotic use and these participant characteristics, they likely reflect known associations between increased infection risk or severity and the need for prolonged antibiotic administration^30–34^. Nevertheless, as potential confounders of the effects of antibiotics on microbiome composition, these variables were included in our subsequent analyses.

**Fig. 1:**
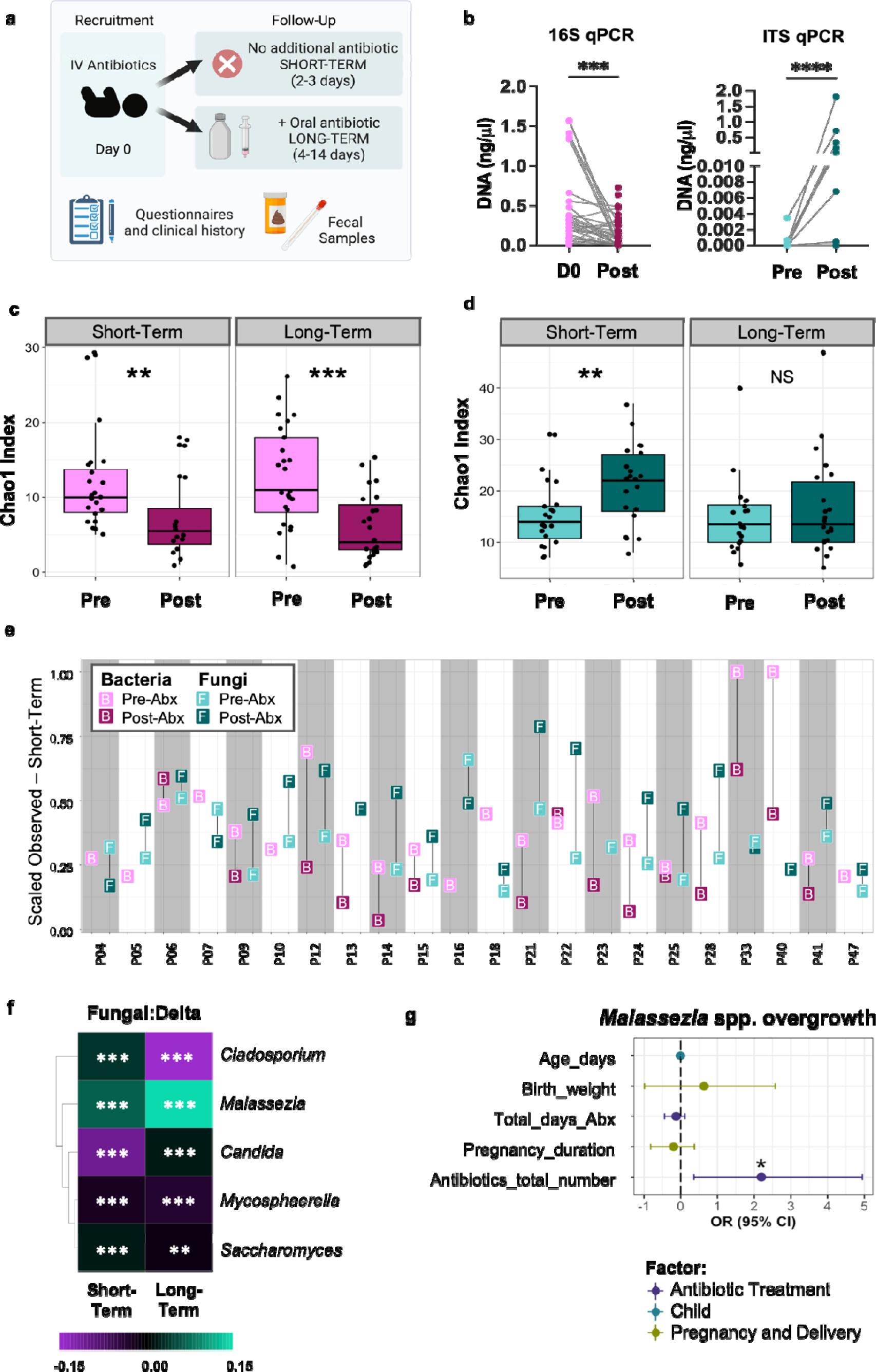
Antibiotic use induces mycobiome alteration in ANTIBIO infants. **(a)** ANTIBIO study design. All participants (< 6 months of age) received intravenous (IV) or intramuscular antibiotics upon recruitment. Following testing and confirmation of infection or not, participants were discharged home with or without a prescription for additional oral antibiotic treatment. Patients were placed into two groups based on the duration of antibiotic treatment: short-term (2-3 days; *n*=22) and long-term (>3 days; *n=*25). Pre-antibiotic samples were obtained within 6 hours from treatment start. Post-antibiotic samples were obtained within 24 hours of cessation of antibiotic treatment. Questionnaires for populational and clinical history were obtained at recruitment and follow-up calls. **(b)** 16S and ITS qPCR quantification (standard curve method) of bacterial and fungal biomass, respectively, pre- and post-antibiotic treatment from the whole cohort of samples. **(c)** Chao1 index plots of bacterial species richness. **(d)** Chao1 index plots of fungal species richness. **(e)** Scaled observed diversity for bacterial and fungal assignments in the short-term treatment group per patient. Values for each participant enrolled in the study are indicated by alternated shading color. **(f)** Delta relative abundance changes for the top 5 most abundant fungal genera identified in ITS2 dataset per study arm. Color denotes increase (green) and decrease (purple) abundance following antibiotic treatment. (c-d) Boxplots indicate median (inside line) and 25th and 75th percentile as the lower and upper hinges, respectively. (b-d, f) Statistically significant differences in relative abundance (pre- vs. post-antibiotics) defined by paired sample Wilcoxon test: \*\**P*<0.01, \*\*\**P*<0.001, \*\*\*\**P<*0.0001. **(g)** Factors associated with *Malassezia* expansion pattern evaluated by multivariable logistic regression. Data presented as log-transformed odds ratio (OR) and 95% confidence interval (CI) for each factor. Expansion: *n*=23, no-expansion: *n*=13. The no-expansion pattern in *Malassezia* abundance was set as the reference level. **\****P*<0.05. Abbreviations: Abx, antibiotic treatment; ER, emergency department.

### Antibiotic use induced strong fungal and bacterial microbiome changes and expansion of *Malassezia* spp. in infants

To explore the effect of antibiotics on the intestinal fungal and bacterial biomass, we determined microbial DNA concentrations in feces by qPCR of the internal transcriber spacer (ITS) region ^35^ and the prokaryotic small subunit ribosomal RNA (16S rRNA) gene ^36^. Antibiotics reduced bacterial and increased fungal DNA concentrations in the complete cohort of samples (**Fig. 1b**). However, antibiotic treatment decreased bacterial concentration only in the short-term but not the long-term group **(Fig. S1a)**, suggesting a possible overgrowth of antibiotic-resistant strains that overtake the newly available ecological niche. In contrast, fungal load was significantly increased after both short- and long-term antibiotic administration **(Fig. S1b)**, indicating that the antibiotic effect on gut fungal biomass was independent of treatment duration.

We then characterized the impact of short- and long-term antibiotic treatment on the intestinal bacterial and fungal microbiomes via shallow shotgun sequencing and analysis using MetaPhAn4.0 ^37^ and ITS2 amplicon sequencing, respectively. Antibiotic treatment had an opposite effect on bacterial and fungal microbiome alpha-diversity. While antibiotics significantly decreased bacterial community species richness (Chao1 index; **Fig. 1c**) and alpha diversity (Shannon index; **Fig. S1c**) in the short- and long-term group, they significantly increased fungal richness (Chao 1; **Fig. 1d**) and alpha-diversity in the short-term treatment group (Shannon index; **Fig. S1d**). To further scrutinize whether the opposing effects of antibiotic use had occurred in the same samples, we examined these effects at the individual level. For the majority of the study participants, antibiotic treatment concomitantly increased fungal alpha-diversity while reducing bacterial alpha-diversity. Overall, antibiotic use increased fungal diversity in 78.9% (15/19) of short-term (**Fig. 1e**), and 56.2% (9/16) of long-term samples **(Fig. S2)**, while reducing bacterial alpha-diversity in 86.7% (13/15) of the short-term (**Fig. 1e**), and 90.9% (20/22) of the long-term samples **(Fig. S2)**. These consistent findings are likely a result of competitive inter-domain interactions due to the disruption of bacterial colonization resistance by antibiotics.

Permutation multivariate analysis of variance (PERMANOVA) of bacterial and fungal community beta-diversity based on Bray-Curtis dissimilarities revealed that antibiotic use, strongly impacted bacterial beta-diversity, especially in the long-term group **(Fig. S1e, Table S2)**. Antibiotic treatment duration significantly explained bacterial community variation in the short-term group (days on antibiotic, R^2^=5.61, *P=*0.02). Other variables significantly associated with bacterial microbiome composition include patient sex, age (days), and birth weight **(Table S2)**. In contrast, antibiotic treatment had little impact on the overall mycobiome community structure of both short- or long-term groups, as well as the entire dataset **(Fig. S1f, Table S2)**. PERMANOVA results showed no significant differences between treatment groups, time points, or duration of antibiotic treatment for fungal communities. Interestingly, in the long-term group, birth weight (R^2^=3.13%; *P=*0.004) and patient sex (R^2^=2.86%; *P=*0.045) significantly explained variance in mycobiome beta-diversity.

We then evaluated the differential abundance of fungal and bacterial taxa after antibiotic treatment (delta variance). Despite the high inter-individual variability in mycobiome composition **(Fig. S3a, c**) median relative abundance values for the top 100 fungal amplicon sequencing variants (ASV) merged at the genus level revealed significant reductions of *Candida, Mycosphaerella, Saccharomyces,* and *Cladosporium*, as well as a notable increase in *Malassezia* in both short- and long-term groups (**Fig. 1f, Fig. S3b, d)**. We further corroborated the results using DESeq2, which revealed that both short- and long-term antibiotic treatment increased the relative abundance of 14 fungal ASVs, with the largest fold changes (8.62 to 26.04 log2 fold change; FC) attributed to three *Malassezia* species: *M. restricta, M. globosa,* and *M. sympodialis* in both treatment groups **(Fig. S3e, Table S3)**. Other fungal ASVs increased with antibiotic use included *Hyphodontia pallidula, Rigidoporus sanguinolentus, Resinicium bicolor,* and *Fomitopsis pinicola* **(Fig. S3e, Table S3)**, all of which are common environmental fungi ^38–42^.

To further examine if the *Malassezia* spp. expansion could be linked to other study variables beyond antibiotic treatment, we used multivariable linear logistic regression analyses on study participant characteristics and antibiotics used in the clinic and/or at home. The associations between study variables and the observed *Malassezia* expansion were adjusted for the confounding effects of age, birth weight, total duration of antibiotic treatment, pregnancy duration, and total number of antibiotics prescribed. Total number of antibiotics received was the only variable positively and significantly associated with the increase in *Malassezia* spp. abundance (OR: 2.1954, CI: 0.3604-4.9453, *P=*0.045; **Fig. 1g, Table S4**).

As expected, infant antibiotic use resulted in very strong compositional changes to the bacterial microbiome. Despite high inter-individual variability **(Fig. S4a-d)**, delta variance analysis identified significant changes in all the 10 most abundant bacterial genera **(Fig. S4e)**. Antibiotic treatment increased the relative abundance of *Staphylococcus* and *Enterococcus,* while reducing *Bifidobacterium, Escherichia, Clostridium,* and *Phocaeicola* genera in both short- and long-term groups **(Fig. S4e)**. DESeq2 further revealed that while most of the differentially abundant bacterial ASVs were reduced, there were a few taxa that increased instead **(Fig. S5)**. These include species of known infectious potential, such as *Klebsiella* spp., *Staphylococcus aureus, Enterococcus avium, Clostridioides difficile,* and *Peptoniphilus harei* ^43–48^. Overall, antibiotics had a striking acute effect on the diversity and composition of the infant gut bacterial and fungal microbiome.

### Antibiotic use impacts microbiome interkingdom interactions

Bacterial-fungal interkingdom interactions are reported in many ecosystems, including the neonatal and infant microbiome ^11,29,49^, yet how antibiotics influence these interactions is less unknown. To this aim, we compared co-occurrence microbial networks at the species level using the NetCoMi (<u>Net</u>work <u>Co</u>nstruction and comparison for <u>Mi</u>crobiome data) before and after antibiotic treatment^50^. Network analysis highlighted interdomain interactions in the infant gut microbiome that included fungi and bacteria **(Fig. S6)**. Before antibiotics, predicted taxa with increased connectivity and positional importance within the network (hub taxa) included the bacteria *Bilophila wadsworthia, Bifidobacterium pseudocatenulatum, Bacteroides ovatus, Clostridium symbiosum,* and *Sutterella wadsworthensis* and the fungi *Mycena arcangelia* and *Hypochnicium geogenium* **(Fig. S6)**. Antibiotic treatment changed the associations and hub taxa within post-antibiotic networks, including the loss of core infant microbiome bacterial taxa *Bifidobacterium* and *Bacteroides*, replaced by opportunistic pathogens *Cutibacterium* spp.*, Prevotella buccalis, Anaerococcus* spp., and *Peptoniphillus lacydonesis* **(Fig. S6)**. Network association analysis revealed differences in the structure of networks before and after antibiotics, with significant differences in measurements for closeness (proximity of nodes; *j=*0.208, *P=*0.03) and eigenvector centrality (central connectedness of nodes within the network; *j=*0.208, *P=*0.03) **(Table S5)**, suggesting potentially different connectivity and interactions within the microbiome. Stark differences in hub taxa between both networks were evidenced in a Jaccard’s index of 0.0 (*P=*0.003), indicating differences in the positional importance of microbial taxa within the networks **(Table S5)**. While network analyses may or may not reflect real biological interactions due to their correlative, hypothesis-generating nature, these data suggest that infant antibiotic treatment impacts interkingdom co-occurrence dynamics in the microbiome. Interestingly, members of the *Malassezia* genus (noted by a star in the networks) displayed increased node sizes, closer proximity, and increased associations with hub taxa after antibiotic treatment **(Fig. S6)**, potentially suggesting increased capacity and importance of microbial interactions. Together, these findings reflect that antibiotic-induced alterations to bacterial and fungal community composition may lead to ecological niche vacancy of core taxa and subsequent occupation and expansion by opportunistic members of the microbiome.

### *M. restricta* colonization shifts the bacterial microbiome in gnotobiotic mice

Species from the *Malassezia* genus have been shown to induce proinflammatory Th1 and Th17 responses in the skin and the gut ^51,52^. Given this and that *M. restricta* was one of the species with the strongest expansion following antibiotic treatment in infants in the ANTIBIO study, we evaluated the effect of colonization with *M. restricta* in gnotobiotic mice. Adult germ-free mice were colonized with defined microbial mixtures containing Oligo-MM12 bacteria (B), or bacteria plus *M. restricta* (B+M) **(Fig. S7a)**. The Oligo-MM12 is a synthetic, stable, inheritable, moderate complexity bacterial consortium ^53,54^, that served as a basis for microbiome-elicited homeostatic immune responses. After breeding, we assessed *M. restricta* colonization in offspring (F1) mice by ITS qPCR in whole colon luminal content. *M. restricta* colonized mice at low amounts throughout neonatal development, even after mice were treated with amoxicillin-clavulanate (t-test, *P=*0.0822; **Fig. S7b-c)**, indicating antibiotic treatment did not impact *M. restricta* colonization in this model.

To explore the impact of *M. restricta* colonization on the coexisting bacterial microbiome, we performed 16S rRNA amplicon sequencing from DNA extracted from fecal pellets of F1 mice 21 days after birth. Permutational analysis on bacterial community beta-diversity (Bray-Curtis) showed that *M. restricta* induced strong community shifts independently of cage effects (colonization, R^2^=33.35, *P*=0.001; **Fig. 2a**). *M. restricta* impacted the relative abundance of key members of the Oligo-MM12 consortium, including an increase in *E. faecalis* (KB1 strain), and a reduction in *Turicimonas muris* (YL45), *Blautia pseudococcoides* (YL58), and *Limosilactobacillus reuteri* (I49) relative abundance (**Fig. 2b-c**). We then used untargeted metabolomics in fecal samples from 21-day-old F1 mice to evaluate the effect of *M. restricta* on microbiome function. Principal component analysis (PCA) plots of the normalized values of 122 characterized metabolites showed substantial shifts in the community metabolic landscape induced by the yeast (**Fig. 2d-e**). Volcano plot analysis yielded seven significantly differentially detected metabolites between B and B+M mice (FC>2, *P* < 0.05; **Fig. 2f, Table S6)**. The most striking change observed in these mice was a 6.56 FC reduction in oleate **(Table S6)**, a fatty acid with a known anti-inflammatory role in mice and humans ^55,56^. *M. restricta*-colonized mice also showed a drastic reduction in cortisol (5.29 FC) and corticosterone (2.23 FC) levels **(Table S6)**, glucocorticoid hormones with broad effects on metabolism, growth, and in regulating homeostatic and immune functions ^57–60^. Overall, these data show that even if detectable at very low amounts, *M. restricta* significantly shifted microbiome composition and functionality.

**Fig. 2:**
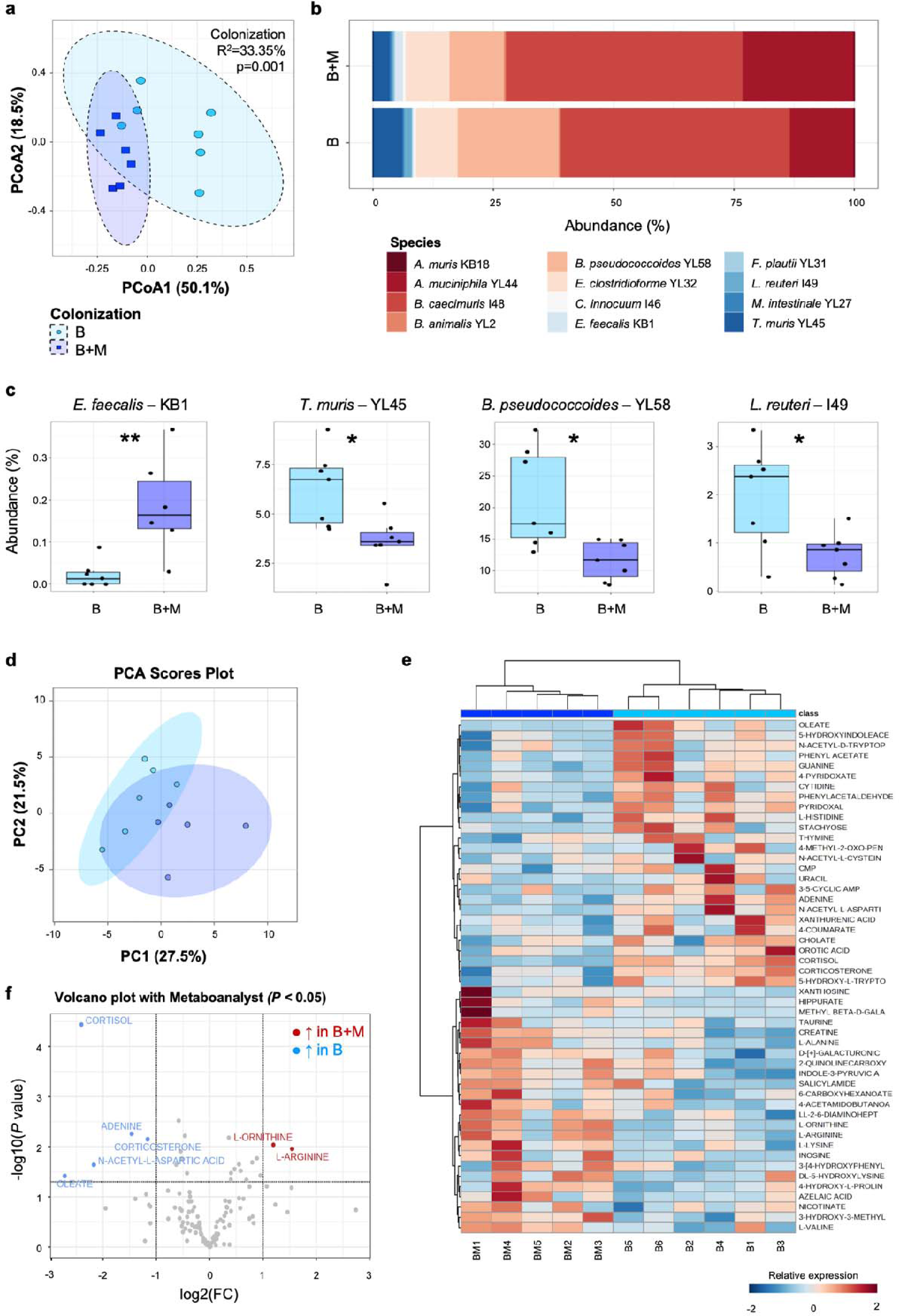
Early-life *M. restricta* exposure shifts microbiome composition and function in gnotobiotic mice. Germ-free mice were colonized with Oligo-MM12 with and without *M. restricta* from birth. **(a)** PCoA plots of bacteria beta-diversity on Bray-Curtis dissimilarities per colonization group. Effect of colonization on community composition defined by PERMANOVA. **(b)** Median relative abundance changes for the 12 members of the Oligo-MM12 consortium. **(c)** Significant changes in bacterial relative abundance. Boxplots indicate the median (inside line) and 25th and 75th percentile as the lower and upper hinges, respectively. Statistically significant differences determined with T-test (YL58, I49, YL45) or Wilcoxon test (KB1), **\****P*<0.05, **\*\****P*<0.01. **(d)** PCA plot of 122 metabolites characterized in fecal samples of gnotobiotic mice at three weeks of age. **(e)** Heat map of top 50 most abundant metabolites in fecal samples. **(f)** Volcano plot with differentially detected metabolites between colonization groups. In red the metabolites increased in B+M mice, in blue the ones increased in B-only mice. Statistically significant differences were determined with FC >2 and T-test, *P*<0.05. Color denotes colonization (BL=Llight blue, B+ML=Lroyal blue). All data points represent biological replicates (sequencing: *N*_B_=7, *N*_B+M_=7; metabolomics: *N*_B_=6, *N*_B+M_=5).

### *M. restricta* exposure induces early-life immune dysregulation

Considering the known role of the microbiome in regulating host immunity, we determined the impact of *M. restricta* colonization on early-life immune functions in F1 mice at 21 days of age. Flow cytometry analysis on immune cells isolated from the colonic lamina propria (cLP) and mesenteric lymph nodes (mLN) showed that *M. restricta* induced strong changes to the intestinal immune landscape. Mice in the B+M group had increased counts of eosinophils, CD64^+^ macrophages, and CD64^+^ cells producing TNF-α (**Fig. 3a-c**), supporting increased macrophage activation. We also detected increased monocyte turnover in the gut, with *M. restricta* colonized mice displaying increased numbers of Ly6C^+^CD64^+^ cells, accounting for both recruited (P1, Ly6C^+^MHC-II^−^) and transitioning (P2, Ly6C^+^MHC-II^+^) monocytes (**Fig. 3d**). Increased monocytic infiltration is a hallmark of cell recruitment to inflammatory sites ^61^, which was also accompanied by increased neutrophil counts in B+M compared to B-only mice (**Fig. 3e**). In addition to these changes in myeloid cells, we also observed increased counts of Th17 and Th2 cells in the cLP of B+M mice (**Fig. 3f-g**), a hallmark of antifungal adaptive immune response and type 2 immunity, respectively ^16,62–66^.

**Fig. 3:**
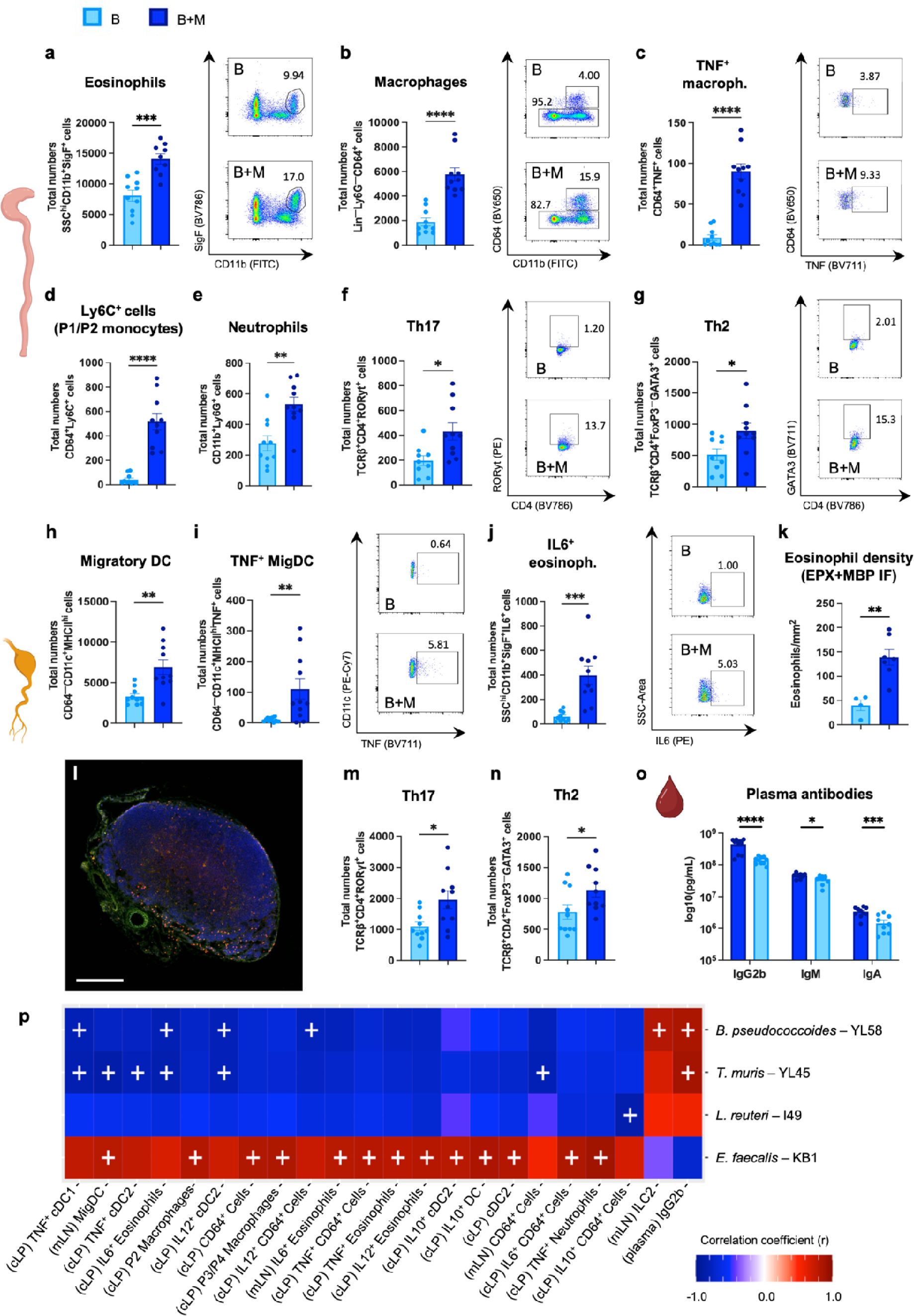
Early-life *M. restricta* colonization induces immune changes in gnotobiotic mice. Immune cell populations and cytokine production were quantified in cLP (a-g) and mLN (h-n) of three-week-old F1 mice. Total numbers of **(a)** eosinophils (plus representative gating for eosinophils), **(b)** CD64^+^ macrophages (plus representative gating for macrophages), **(c)** TNF^+^ macrophages (plus representative TNF staining in macrophage population), **(d)** Ly6C^+^ monocytes, **(e)** neutrophils, **(f)** Th17 (plus representative RORγt staining in CD4^+^ population), and **(g)** Th2 cells (plus representative GATA3 staining in CD4^+^FoxP3^−^ population) in cLP. Total numbers of **(h)** MigDC, **(i)** TNF^+^ MigDC (plus representative TNF staining in MigDC population), and **(j)** IL-6+ eosinophils (plus representative IL-6 staining in eosinophil population). **(k)** Normalized counts of eosinophils in mLN quantified using IF. **(l)** Representative IF image of 21-day-old mLN of B+M mice. Sectioned slides were stained with anti-MBP (AF647; red), anti-EPX (AF488; green), and DAPI (blue). Eosinophils were characterized by overlay of MPB and EPX signal. Scale bar = 200 μm. Total numbers of **(m)** Th17 and **(n)** Th2 cells in mLN. **(o)** Antibodies determined in plasma using Mouse Isotyping Panel 1 kit. (a-o) Color denotes colonization treatment (BL=Llight blue, B+ML=Lroyal blue). Data represented as mean ± S.E.M. All data points represent biological replicates (cLP, mLN, plasma, *N*_B_=10, *N*_B+M_=10; IF, *N_B_*=4, *N_B+M_*=8). Statistically significant differences defined by student unpaired T-test; \**P*<0.05, \*\**P* <0.01, \*\*\**P* <0.001, \*\*\*\**P* <0.0001. **(p)** Heat map of biweight correlations between merged bacterial ASV at the species level (x-axis) and reported immune features (y-axis) at day 21. Color denotes positive (red) and negative (blue) correlation values. Significant correlations are denoted with a cross and defined by the BiCOR method with FDR correction; *P*L<L0.05.

We also determined the impact of *M. restricta* colonization on immune responses in the draining mLN, as this is an important immune priming site for both intestinal and systemic immune responses ^67,68^. B+M mice displayed increased numbers of migratory dendritic cells (MigDC; CD11c^+^MHC-II^hi^) and MigDCs producing TNF-α in the mLN (**Fig. 3h-i**), potentially indicating higher fungal antigen trafficking for induction of adaptive immunity at this site. Eosinophils producing IL-6, known to promote Th2 differentiation ^69^, were also elevated in B+M mice (**Fig. 3j**). By labelling and quantifying eosinophils with immunofluorescence (IF) microscopy, we confirmed that *M. restricta* increased eosinophil counts in mLN, and revealed eosinophils accumulated in the paracortical region on this lymphoid organ (**Fig. 3k-l**), where T and B cell regions are localized ^70,71^. Like in the cLP, we also observed increased Th2 and Th17 cells in the mLN of mice colonized with *M. restricta* (**Fig. 3m-n**). Several other immune cell types were not impacted by *M. restricta* colonization in cLP and mLN, including Th1 cells, Tregs, CD4^+^ and CD8^+^ T cells (**Fig S8**). Finally, to also assess the impact of *M. restricta* on systemic humoral immune responses, we determined antibody titers in the plasma of F1 mice. Mice exposed to *M. restricta* showed reduced concentrations of IgG2b, IgA, and IgM plasma antibodies (**Fig. 3o**), supporting a generalized effect on B cell response in B+M group. Altogether, these data support that *Malassezia*-derived signals alter early-life immune responses in the gut and draining mLN of gnotobiotic mice, including cell types deemed critical in atopy development.

To evaluate the potential associations between the described immune changes and the microbiome, we correlated the relative abundance of the bacterial species (**Fig. 2c**) with immune features in plasma, cLP, and mLN (**Fig. 3**) using the Biconjugate A-Orthogonal Residual (BiCOR) method ^72^. This analysis yielded significant associations between immune changes and bacterial taxa differentially impacted by *M. restricta* colonization (**Fig. 3p**). Notably, *E. faecalis* (KB1), the only bacterial taxon increased in B+M mice, had the largest number of significant and mostly positive associations with several immune readouts, including a positive correlation with CD64^+^ macrophages, P2 and P3/P4 (CD64^+^Ly6C^−^) macrophage subtypes, type-2 conventional DC (cDC2; CD11b^+^), MigDC, and IL-6, IL-10, IL-12, and TNF-α cytokine production by different cell types (**Fig. 3p**). This suggests that the immune effects observed may be linked to changes in this bacterial species, which is known to colonize the murine gut close to the epithelium ^73^, potentially increasing local immune responses during dysbiosis. In contrast, *B. pseudococcoides* (YL58), *T. muris* (YL45) and *L. reuteri* (I49), all of which decreased in abundance upon *M. restricta* exposure, were inversely correlated with some of the myeloid immune cell types found elevated in B+M mice (**Fig. 3p**).

While *M. restricta* exerted strong immunomodulatory effects in gnotobiotic mice, we sought to determine if this was due to working with mice colonized with a low-diversity bacterial community composed mostly of low and medium immunogenicity species. Thus, to differentiate *M. restricta*-induced immune responses from other immunogenic gut commensals, we upgraded the Oligo-MM12 consortium to also include *C. albicans* (isolate K3) and *Pseudomonas fluorescens* (isolate J4) (B+CP), isolated from ANTIBIO infant stool samples, and compared it to mice colonized with all species and *M. restricta* (B+CP+M). *C. albicans* is a common intestinal colonizer, identified in fecal samples from a very early age ^74,75^, and induces strong mucosal immune responses ^16,62,76^. *P. fluorescens* is a recently appreciated member of the intestinal microbiome, with strong immunogenic effects in mammals ^77,78^. We observed very similar immune effects when *M. restricta* exposure occurred in combination with *C. albicans* and *P. fluorescens* (B+CP+M), such as increased eosinophils, macrophages, Th2, and Th17 cell counts in cLP, compared to B+CP mice **(Fig. S9a)**. We also detected a trend in increased TNF-α^+^ MigDC, and a significant increase in Th2 and Th17 counts in mLN of B+CP+M mice compared to B+CP **(Fig. S9b)**. Altogether, these data show that *M. restricta* uniquely induces local and systemic early-life immune dysregulation, even in mice colonized with other immunogenic bacterial or fungal species.

### Early-life *M. restricta* colonization enhances airway inflammation in mice

To determine the causal role of early-life *M. restricta* colonization and induced immune dysregulation on allergic airway disease, F1 gnotobiotic mice were subjected to three weekly intranasal HDM challenges at 7-10 weeks of age (**Fig. 4a**). Using this model, only B+M mice showed overt airway inflammation, marked by increased inflammatory infiltration in their bronchoalveolar lavage fluid (BALF) and marked BALF eosinophilia, compared to unchallenged, naive controls (**Fig. 4b-d**). Histopathological scoring showed that *M. restricta* colonization significantly increased lung inflammation, accounting for perivascular, peribronchial, and parenchymal infiltration, as well as epithelial damage, compared to naive and B-only mice (**Fig. 4e-f**). These data further support the unique effects of early-life *M. restricta* on not only gut immune dysregulation, but also increasing susceptibility to allergic airway inflammation against common aeroallergens.

**Fig. 4:**
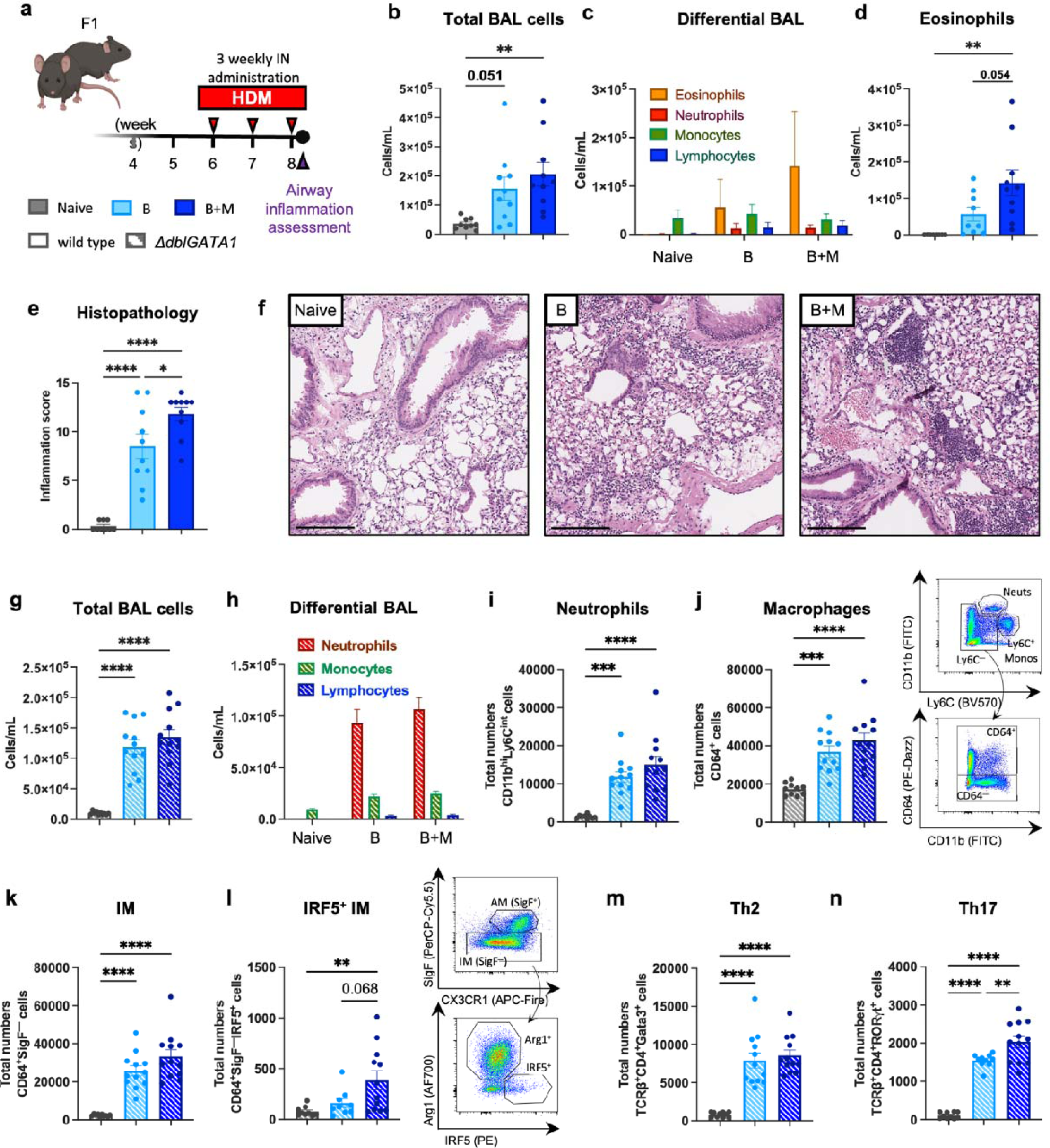
Early-life *M. restricta* exposure increases allergic airway inflammation. **(a)** Experimental layout for HDM airway inflammation model. F1 mice in the B and B+M groups were intranasally immunized with three HDM challenges at weeks 6, 7 and 8 post-birth. 24h after the last challenge, inflammatory response was assessed and compared to naive mice that received PBS as a control. **(b)** Total cell counts in BALF of wild-type mice. **(c)** Compiled differential cell counts of BALF infiltrate. **(d)** Total eosinophil counts in BALF. **(e)** Histopathological scores of challenged airways. **(f)** Representative hematoxylin and eosin-stained sections of challenged airways of wild-type mice. Scale bar = 200 μm. **(g)** Total cell counts in BALF of Δ*dblGATA1* mice. **(h)** Compiled differential cell counts of BALF infiltrate. Total numbers of **(i)** neutrophils, **(j)** CD64**^+^** macrophages (plus representative gating for neutrophils and macrophages), **(k)** IM (CD64^+^SigF^−^), **(l)** IRF5^+^ IM (plus representative IM and IRF5 staining in IM population), **(m)** Th2, and **(n)** Th17 cells in lungs of Δ*dblGATA1* mice. Color denotes colonization treatment (Naive = gray, BL=Llight blue, B+ML=Lroyal blue), and strike-out effect denotes genetic background (wild type= solid, Δ*dblGATA1*=strike out). Data represented as mean ± S.E.M. All data points represent biological replicates (wild-type, *N*_naive_=9, *N*_B_=10, *N*_B+M_=10; Δ*dblGATA1, N*_naive_=10, *N*_B_=12, *N*_B+M_=13). Statistically significant differences defined by ANOVA + Tukey post-hoc correction; *p<0.05, **p<0.01, ***p<0.001, ****p<0.0001.

### *M. restricta*-induced immune dysregulation and airway inflammation are partially dependent on eosinophils

A consistent observation across our *M. restricta* colonization model was the significant eosinophilia in the colon, mLN, and lungs. Eosinophils are tissue-resident immune cells that support intestinal homeostasis in response to microbiota signals ^79^. In addition to their ability to exert antifungal activity, eosinophils mediate and support immune responses in the host via recruitment and activation of other immune cells ^79–82^. To determine if *Malassezia*-induced eosinophilia orchestrates the observed immune dysregulation and airway inflammation, we repeated the colonization and HDM inflammation models in eosinophil-deficient Δ*dblGATA1* mice ^83^. *M. restricta* colonization in Δ*dblGATA1* mice ablated the *M. restricta* effect on macrophages, dendritic cells, and myeloid cytokine responses in cLP and mLN seen in wild-type mice colonized with this fungus, yet still induced increased neutrophil counts in their cLP and Th2 and Th17 cells in both cLP and mLN **(Fig. S10a-b)**. This experiment revealed that *M. restricta*-induced myeloid immune responses are eosinophil dependent, while the Th-skewed immune phenotype is likely mediated by other intestinal immune cells.

To determine if eosinophils were required for *M. restricta* immune responses in the airways, we subjected Δ*dblGATA1* mice to the same HDM challenge scheme as wild-type mice (**Fig. 4a**). In agreement with previous reports showing eosinophils are dispensable for certain aspects of inflammatory responses to allergens ^84–86^, Δ*dblGATA1* mice developed robust airway inflammation (**Fig. 4g**), with marked infiltration of neutrophils and macrophage/monocytes (**Fig. 4h**). Additional characterization of the immune response in the lungs using flow cytometry showed that most of the cell types assessed were present in similar counts between B and B+M Δ*dblGATA1* mice, although there was an increase in IFN regulatory factor 5 (IRF5)-expressing interstitial macrophages (IM, CD64^+^SigF^−^) and Th17 cells in Δ*dblGATA1* B+M mice (**Fig. 4i-n**), further implicating eosinophils in the *M. restricta*-induced immune responses to aeroallergens in the lungs.

## DISCUSSION

It is well established that antibiotic-induced intestinal microbiome dysbiosis predisposes children to develop asthma ^20–24^ and increases the severity of allergic inflammation in murine models ^25–28,87^. We have also previously shown that intestinal fungal colonization exacerbated subsequent airway inflammation to ovalbumin allergen in gnotobiotic mice ^29^. The work presented in this study expands on this by distilling the complex interactions between the largely unexplored fungal dysbiosis, immune dysregulation and susceptibility to allergic airway inflammation. Our study corroborated the powerful effect of infant antibiotic treatment on the bacterial gut microbiome, with dozens of species impacted, yet it also showed how the mycobiome responds to this strong ecological perturbance. Even parenteral antibiotic courses of less than 72 hours resulted in an increase in fungal abundance and species diversity explained by fungal taxa that benefitted from a transient loss in bacterial colonization resistance. Most prominently, species from the *Malassezia* genus expanded in the infant gut during both short- and long-term antibiotic treatment (**Fig. 1f, Fig. S3**). Colonization of gnotobiotic mice with *M. restricta,* one of the fungal species most heavily favored by antibiotic use, resulted in pronounced shifts in bacterial microbiome composition and function (**Fig. 2**). *M. restricta* colonization also induced marked changes in the early-life intestinal immune landscape, including increased counts of eosinophils, macrophages, Th2, and Th17 cells (**Fig. 3**), indicating immune dysregulation that rendered these mice more susceptible to allergic airway inflammation (**Fig. 4**).

To our knowledge, the ANTIBIO Study is the first characterization of the acute effect of antibiotic use on the infant mycobiome. Although this study lacked an age-matched untreated control group, it was designed with sufficient power to detect an effect on the mycobiome based on previous infant studies^9^. Further, this study enabled the sequencing-based characterization of the acute effects of antibiotic administration on microbiome diversity and composition at the individual level. In most infants, even those receiving only parenteral therapy, antibiotics had opposing effects on bacterial and fungal alpha-diversities, modifying cross-kingdom interactions and overall network characteristics. Our analysis also revealed that, while antibiotic use exerted the strongest effect on community diversity and composition, other factors, such as patient’s age, sex, and birth weight also impacted the microbiome. These may have influenced how the microbiome responded to antibiotics, as seen in certain study participants who exhibited atypical changes in microbiome diversity after antibiotic use (i.e.: a decrease in fungal diversity and/or an increase in bacterial diversity) (**Fig. 1e, Fig. S2)**. Thus, the complex interactions between clinical and environmental factors known to impact the bacterial microbiome are also present for the mycobiome.

While the ANTIBIO study was designed to detect microbiome alterations associated with antibiotic treatment (primary outcome), our study also showed that combination antibiotic therapy was significantly associated with *Malassezia* spp. expansion after controlling for confounding factors, such as gestational age at birth, antibiotic therapy duration, and birth weight (**Fig 1g**). In line with this, reduced birth weight and antibiotic treatment duration have been associated with *Malassezia* skin colonization, and lung and systemic infections in term and preterm infants^88–91^. Species of this genus, such as *M. globosa, M. restricta, M. sympodialis,* and *M. furfur* are common colonizers of the skin ^92,93^ but some are commonly found in the gut as well, particularly during infancy ^8,75,94,95^. Patterns of mycobiome development during the first year of life reveal a higher predominance of *Malassezia* spp. that is later replaced by Saccharomycetales and *Candida*, possibly as a response of the introduction of complimentary feeding after 6 months of age. All species within the *Malassezia* genus are unable to synthesize fatty acids or have unique lipid dependencies requiring external sources of lipids for growth ^96^. It is possible that the higher lipid content in breast milk and formula supports better growth conditions for these species in the gut during the period of exclusive lactation. Supporting this, we found a reduction in oleic acid concentration, a fatty acid found in the highest concentration in breast milk^97^, in young mice colonized with *M. restricta* (**Fig 2f**).

We chose to center our efforts on *M. restricta,* given the pronounced effect of antibiotic use on its abundance and its previously reported role in exacerbating chemically-induced colitis in mice ^52^. While we encountered differences in colonization ability between infants and mice (as it often occurs in human microbiota-associated mice^98^), *M. restricta* was detected during the first three weeks of life, even 16 days after exposure **(Fig. S7),** suggesting its ability to colonize the mouse colon. Even in low amounts, *M. restricta* induced significant taxonomic and functional changes in the mouse microbiome, indicating that this fungal species interacts with bacterial members of the microbiome, as has been previously reported in the skin ^99,100^. This is also supported by our network analysis of the gut microbiome in 3- and 12-month-old Canadian infants, showing *Malassezia* species as hub taxa predicted to have positional importance within the community^95^. The ability of *Malassezia* spp. to modulate microbiome composition may also suggest that its expansion during antibiotic treatment may contribute or even exacerbate intestinal microbiome dysbiosis via interkingdom interactions.

The immunomodulatory role of *Malassezia* spp. has been previously reported in skin and gut inflammatory conditions ^51,52,63,101^. Interestingly, *Malassezia* spp. emerged as the fungal taxa most strongly associated with the *CARD9^S12N^* risk allele for inflammatory bowel disease ^52^. This study additionally showed that *M. restricta* colonization exacerbated DSS-induced colitis in mice, via innate CARD9 signaling ^52^. Our data further reveals that colonization with this yeast induces broad immune dysregulation independently of chemically-induced inflammation or infection. Even at low levels, colonization with this yeast resulted in long-lasting and widespread immune effects, such as increased intestinal pro-inflammatory infiltrate and skewed Th2 and Th17 responses, reduced systemic antibody production, and increased susceptibility to allergic airway inflammation later in life. Importantly, our work suggests that these effects are unique for this species, as other immunogenic microbes, such as *C. albicans* or *P. fluorescens* did not elicit the same immune responses.

Our experimental design focused on ecological and immune mechanisms impacted by *M. restricta*, thus microbial-derived mechanisms were not directly investigated. It is possible that this yeast induces direct effects by interacting with intestinal immune cells, as *Malassezia* can be found in close association with the intestinal mucosa ^52^, where it can be directly sensed and taken up by phagocytes ^76,102^. Simultaneously or alternatively, *Malassezia* may exert its immune effects indirectly via changes to the gut microbiome. For instance, our metabolomics results showed marked reductions in immunoregulatory metabolites with known links to microbial metabolism, such as oleate, cortisol, and corticosterone ^55,56,103,104^. Oleic acid is a known anti-inflammatory fatty acid that modulates immune responses in both innate and adaptive systems ^55,56^, with protective effects in colitis models ^105,106^. Considering its lipid-dependent nature ^96^, *M. restricta* could be directly consuming oleate, reducing its absorption and potential anti-inflammatory roles. Further, the levels of glucocorticoid hormones cortisol and corticosterone change during development ^107,108^, are influenced by microbiota composition ^109,110^, and are modulated by fungal exposure ^111,112^. Glucocorticoids act on the intracellular glucocorticoid receptor to regulate the transcription of genes in leukocytes, influencing Th cell development ^103^ and myeloid cell functions ^104^. While not directly explored in our work, it is possible that metabolic shifts induced by colonization with *M. restricta* exert the immune responses observed.

An important observation of our study is how this fungus induced eosinophilia in several mouse tissues. Eosinophils actively recognize fungal-associated molecular patterns, exert antifungal activity, and mediate immune responses in the host upon fungal exposure ^80^. Eosinophils secrete a myriad of immunoregulatory molecules, including granule pre-formed proteins (e.g.: MBP, EPX), enzymes (e.g.: catalase, phospholipase), and cytokines/chemokines (e.g.: IL-4, IL-5, IL-13, IL-6, IFN-γ, GM-CSF, CCL3, CCL5), which synergistically support immune responses with other cells ^81,82^. For example, eosinophil deficiency (i.e.: *PHIL*, α-IL-5) results in increased colitis severity following DSS treatment ^113,114^. Eosinophils also support immune cell migration and intestinal homeostasis in response to microbiota signals ^79^. Our data from Δ*dblGATA1* gnotobiotic mice show that *Malassezia*-induced immune dysregulation is partially dependent on eosinophils, abrogating myeloid immune responses in *M. restricta*-colonized mice. This is in line with the reported reduced myeloid functions and inflammatory responses in Δ*dblGATA1* mice ^115^, underlining the role of eosinophils in orchestrating myeloid cell recruitment **(Fig. S10)**. Nevertheless, our data show that the absence of eosinophils did not impact Th2 and Th17 cells in cLP, mLN, and HDM-challenged lungs (**Fig. 4, S10).** This is in agreement with a previous publication showing eosinophils are dispensable for intestinal adaptive responses in a *Giardia muris* infection model ^116^, and that immune responses to intestinal fungal colonizers increase Th17 induction in the airways ^13,62,117^.

Despite showing infant antibiotic use strongly impacted the microbiome, and that a fungal taxon expanded in post-antibiotic treated samples causally impacted immune responses in mice, it is important to consider our study limitations. First, much larger, prospective studies are needed to determine if antibiotic-induced *Malassezia* spp. expansion in infants is transient or not, and whether it mediates later risk to atopic disorders, like asthma. Also, while we chose a well-established bacterial consortium, these 12 bacteria are unable to recapitulate the same level of immune response in the host as more complex microbial communities ^118,119^. Future studies should examine the ecological and immune consequences of *Malassezia* spp. colonization in mice harboring more complex microbiomes. Our work also did not investigate if *M. restricta* colonized other tissues in mice, such as the airway or the skin, potentially exerting relevant immune responses at those sites. For example, *Malassezia* has been associated with acute pulmonary exacerbations in cystic fibrosis patients^120^ and has also been found elevated in induced sputum samples of asthmatics compared to health controls ^121^. Finally, additional efforts are needed to identify other immune mechanisms beyond eosinophil function. Ultimately, our reverse-translational approach has characterized a causal role of *M. restricta* in altering intestinal microbiome composition and function, inducing immune dysregulation in the host, and exacerbating airway inflammation in gnotobiotic mice. These data highlight novel mechanisms of mycobiome-induced immune dysregulation originating from common practice antibiotic therapy in early life.

## METHODS

### Study design

This study used a reverse translational approach by combining a population-based, observational, prospective clinical study in young infants receiving antibiotics in the Calgary metropolitan area, Alberta, Canada, followed by experiments in gnotobiotic mice to determine the impact of fungi expanded after antibiotic treatment in microbiome ecology and host immune development. The ANTIBIO clinical study protocols were approved by the Calgary Conjoint Health Research Ethics Board (REB17-1951). The caregivers of all participants provided written informed consent. The gnotobiotic mouse experiments were conducted under protocols approved by the University of Calgary Animal Care Committee (AC20-0176), following the guidelines of the Canadian Council on Animal Care.

### ANTIBIO study recruitment and sample collection

To study the impact of antibiotic use on the infant bacterial and fungal microbiomes, the ANTIBIO study recruited 47 infants aged <6 months of age evaluated in the ACH-ED, in Calgary, Alberta from July 2018 to February 2020. To differentiate the microbiome changes attributed to short- and long-term antibiotic treatment, study participants were categorized into two groups: (i) infants receiving antibiotics for 2-3 days (short-term), or (ii) infants receiving antibiotics for 4-14 days (long-term). Two fecal samples were obtained per study participant. An initial stool sample or rectal swab was collected in the hospital by a research nurse from the Pediatric Emergency Research Team (PERT) within six hours of the first dose of antibiotics (pre-antibiotics). A second stool sample or rectal swab was collected within 24 hours of the last dose of antibiotic (post-antibiotics). The follow-up sample was collected either by a PERT nurse in the hospital, or by the patient’s caregiver if discharged home with a prescription of oral antibiotics. Antibiotic treatment consisted of parenteral antibiotics received in the hospital or the combination of parenteral plus oral antibiotics prescribed at discharge from the ACH-ED. Enrolment questionnaires were completed in person by the caregivers of all participants. Additionally, PERT Research Nurses collected information from charts and medical records regarding admission and treatment outcomes. Follow-up questionnaires were conduced over the phone on the last day of antibiotic treatment by a member of the Arrieta lab (Dr. Marie-Claire Arrieta or Erik van Tilburg Bernardes). The study was conducted in accordance with Good Clinical Practice guidelines and all applicable regulations. Additional ANTIBIO study details are provided in *Supplementary Materials*.

### Microbial DNA isolation and quantification

Microbial DNA was obtained from fecal samples (humans or mice) using the DNeasy PowerSoil Pro kit (Qiagen) according to manufacturer’s instructions with an added cell lysis step at TissueLyser II bead beater (Qiagen), eluted in nuclease-free water, and stored at −80°C. Fungal and bacterial DNA concentrations in fecal samples were determined by qPCR, as previously described ^29,35,36^. The ITS marker and 16S rRNA gene were selected for quantitative amplification of fungal and bacterial DNA, respectively, using iQ SYBR Green Supermix (BioRad Laboratories). The qPCR runs were performed on the StepOne Plus Real-Time PCR System (Applied Biosystems) in the Snyder Resource Laboratories, University of Calgary. Detailed qPCR protocols are described in *Supplementary Materials*.

### Shotgun metagenomics sequencing, processing, and bacterial taxonomic assignment

In-house extracted DNA samples were sent to Microbiome Insights (Vancouver, Canada) for shallow shotgun sequencing. Shotgun sequencing was performed as previously described ^4^, with modifications. Libraries were prepared using an Illumina Nextera library preparation kit (Illumina) and sequenced in a NextSeq 500 System (Illumina) using paired-end (150 bp x 2) reads. Shotgun metagenomic sequence reads were processed in four steps: adapter removal, read trimming, low-complexity-reads removal, and host-sequence removals. The remaining reads were taxonomically classified using MetaPhlAn4.0 ^37^.

### ITS2 and 16S sequencing, processing, and bacterial taxonomic assignment

In-house extracted DNA samples were sent to Microbiome Insights (Vancouver, Canada) for ITS2 and 16S amplicon sequencing. The ITS2 and 16S V4 regions were amplified and sequenced as previously described ^29,75,122^. Demultiplexed forward and reverse reads were denoised, filtered, inferenced, trimmed, and merged using DADA2 v.1.26.0 ^123^ package for R (R Development Core Team; http://www.R-project.org). Samples with less than 1,000 sequencing reads were excluded and ASVs appearing in only one sample (singletons) were removed. Taxonomic assignment of the resulting ASVs was performed using the UNITE database^124^, or an in-house database containing the 16S gene sequences of the Oligo-MM12 consortia strains^125^. The remaining dataset was filtered for ASV appearing at least three times in 10% of samples.

### Gnotobiotic mouse model of *M. restricta* colonization

Germ-free wild-type C57BL/6J mice were obtained from in-house breeding colonies in the International Microbiome Centre (IMC). Specific pathogen-free Δ*dblGATA1* (B6.129S1(C)-Gata1^tm6Sho^/LvtzJ; JAX Stock No. 033551) mice were purchased from The Jackson Laboratory and rederived germ-free through the axenic embryo transfer program in the IMC. Eight- to 15-week-old germ-free wild type or Δ*dblGATA1* C57Bl/6J female mice were orally gavaged with a consortium of microorganisms prepared under anaerobic conditions by mixing 100 μL of microbial cultures, as previously described^29^. The consortia consisted of twelve mouse-derived bacteria species (Oligo-MM12)^54^ with or without: *M. restricta* (ATCC MYA-4611)*, C. albicans* (isolate K3), and/or *Pseudomonas fluorescens* (isolate J4). Oligo-MM12 bacteria were grown for 2 days at 37°C in fastidious anaerobe broth (LabM, Heywood, Lancashire, UK) under anaerobic conditions. *M. restricta* was grown for 7-10 days at 30°C in mDixon broth. *C. albicans* K3 and *P. fluorescens* J4 clinical isolates were grown for 1 day at 30°C in YM broth (BD Biosciences, San Diego, CA, USA). After gavages, mice were paired for mating in a 2:1 female:male ratio per cage. To ensure fungal exposure to F1 mice, corresponding fungal cultures were spread twice on the dam’s abdominal and nipple regions during the pups’ first week of life, as previously described ^3,29^. To gain further insight into the implications of antibiotic use in *M. restricta* colonization, a group of B+M mice was treated with antibiotic amoxicillin-clavulanate (0.2 mg/ml; Sigma, Oakville, ON, Canada; “B+M+Abx”) from the 2^nd^ to 21^st^ day post birth. Mice were housed inside gnotobiotic flexible-film isolators in the IMC, under a 12-h light/12-h dark cycle, 40% relative humidity, 22–25°C, and *ad libitum* access to sterile food and water.

### Metabolomics sample preparation, run, and analysis

Fecal samples were obtained from F1 mice at 21 days of age and processed for metabolomics, as previously described ^29,122^. Samples were subjected to untargeted metabolomics using liquid chromatography-mass spectrometry (LC-MS) at the Calgary Metabolomics Research Facility (CMRF), at the University of Calgary, which is supported by the IMC. Files containing metabolite concentration were received from CMRF, and concentration data were normalized through median, square root transformation, and Pareto scaling on MetaboAnalyst v.5.0 ^126^.

### Early-life immune assessment

Early-life immune functions were determined in mouse plasma, cLP, and mLN at 21 days post-birth. Three-week-old F1 mice were anesthetized with isoflurane. Blood was drawn by cardiac puncture in syringes primed with heparin (100 U/mL; Sandoz Canada Inc., Boucherville, QC, Canada), and mice were euthanized by cervical dislocation. Plasma was separated by centrifugation at 3,000 rcf for 10 min, at 4°C. IgA, IgG1, IgG2a, IgG2b, IgG3, and IgM titers were determined in plasma via electrochemiluminescence, in Mouse Isotyping Panel 1 Assay Kit (Meso Scale Diagnostics, Rockville, MD, USA), as previously described ^29^. The whole colon and mLN were surgically excised for immune cell isolation. The intestinal immune landscape was characterized in cLP and mLN using flow cytometry, as described below.

### Isolation of intestinal immune cells

cLP immune cell isolation was performed as previously described ^79^, with small modifications. Whole colons were excised, cleaned of any fat and connective tissue, cut longitudinally, and washed in ice-cold Hank’s Balanced Salt Solution (HBSS) supplemented with 2% inactivated horse serum (iHS; Sigma). Epithelial layers were removed with two 20 min incubation, at 37°C in pre-warmed epithelial cell removal solution (HBSS supplemented with 4% iHS [Sigma] and 5 mM EDTA [Sigma]). The remaining tissues containing lamina propria and muscularis layer were digested for 20 min digestion, at 37°C in pre-warmed digestion medium (RPMI supplemented with 10 mM HEPES [Thermo-Fisher, Waltham, MA, USA], 10 % iHS [Sigma], 10 U/mL DNase I [Sigma], and 1 mg/mL Collagenase Type VIII [Sigma]). Following digestion, the cells were filtered through a 100 µm cell strainer (Thermo-Fisher), pelleted down, resuspended in RPMI supplemented with 10% iHS, and immediately stained for flow cytometry.

### Isolation of mLN immune cells

mLN immune cell isolation was performed as previously described ^127^, with small modifications. mLN were cleaned of any fat and connective tissue and digested for 25 min, at 37°C in digestion medium (RPMI supplemented with 1 mg/ml Collagenase IA [Sigma] and 30 U/ml DNase I [Sigma]). Following digestion, the cells were filtered through a 100 µm cell strainer (Thermo-Fisher), pelleted down, resuspended in RPMI supplemented with 10% iHS, and immediately stained for flow cytometry.

### HDM airway inflammation model

An experimental murine HDM (*Dermatophagoides pteronyssinus*) model was performed as previously described ^117^. Briefly, seven- to ten-week-old F1 mice were immunized intranasally (IN) with 100 µg HDM (Greer Laboratories, Lenoir, NC, USA) in 50 µL sterile PBS (2 mg/mL) three times with 7-day intervals, under light isoflurane anesthesia. Additionally, a pooled group of mice from different colonization groups received vehicle solution (50 µL sterile PBS) and denominated naive. Airway inflammation was assessed one day after the last IN challenge. BALF retrieval was performed by 3xL1 mL washes with ice-cold PBSL+L10% iHS, in mice anesthetized with ketamine (200 mg/kg; Vetoquinol) and xylazine (10 mg/kg; Bayer Inc.). The recovered BALF was combined in a 15 mL falcon and kept on ice for total and differential cell counts. Total cell counts in BALF were performed in a hemocytometer. Differentials (eosinophils, neutrophils, macrophages, and lymphocytes) were performed from 200 cells in hematoxylin and eosin-stained CytoSpins (Shandon Cytospin II, Thermo-Shandon, Runcorn, Cheshire, UK) based on standard morphological criteria.

### Lung histology

Following BALF retrieval, the left lung was inflated, excised, and fixed in 10% formalin. Lungs were sent to the Alberta Province Laboratories (APL) for paraffin embedding, cut into 4 μm sections, and hematoxylin and eosin staining. Sections were blindly scored, as previously described ^20^, with small modifications. Sections were scored on a 0-to-4 scale (0 = none, 4 = severe disease) for the following parameters: (i) peribronchial infiltration, (ii) perivascular infiltration, (iii) parenchymal infiltration, and (iv) epithelium damage, for a maximum score of 16. Following scoring, representative slides were selected for imaging, using an Aperio AT2 slide scanning microscope (Leica Biosystems, Wetzlar, Germany), at 40 x magnification. Images had dimensions set in QuPath v.0.4.3 bioimage analyzer ^128^, exported in TIFF format, and opened in ImageJ 1.53t (http://imagej.net) to overlay scale bars (200 μm).

### Isolation of lung immune cells

Following BALF retrieval, the right (inferior and post-caval) lobules were excised and collected in RPMI supplemented with 2% iHS. Lung immune cell isolation was performed as previously described ^129^, with modifications. Lobules were minced and digested for 30 min, at 37°C in digestion medium (RPMI supplemented with 2% iHS [Sigma], 1 mg/ml Collagenase XI [Sigma], and 10 U/ml DNase I [Sigma]). Following digestion, the cells were filtered through a 100 µm cell strainer (Thermo-Fisher) and residual red blood cells were lysed in ACK lysis buffer (Quality Biological, Gaithersburg, USA) for 7 min. Cells were pelleted down resuspended in RPMI supplemented with 10% iHS and immediately stained for flow cytometry.

### Immune cell staining and flow cytometry

Isolated immune cells (from cLP, mLN, or lungs) were stained for 20 min at room temperature with eFluor506 Fixable Viability Dye (Invitrogen, Carlsbad, CA, USA). Cells were then incubated in Fcγ receptor blocking antibody (BD Biosciences) and an array of intra and extracellular markers for immune cell characterization. Cells were stained for surface markers for 20 min, fixed and permeabilized with eBioscience Foxp3/Transcription Factor Staining Buffer Set (Invitrogen), according to manufacturer’s protocols, and stained for intracellular markers overnight at 4°C. The following antibodies were used: BV421 anti-T-bet (clone 4B10; BioLegend, Sand Diego, CA, USA), eF450 anti-IL-5 (TRFK5; Invitrogen), BV570 anti-CD8 (53-6.7; BioLegend), BV605 anti-IL-10 (JES5-16E3; BD Biosciences), BV650 anti-IFN-γ (XMG1.2; BD Biosciences), BV711 anti-GATA3 (L50-823; BD Biosciences), BV750 anti-TCRβ (H57-597; BD Biosciences), BV786 anti-CD4 (RM4-5; BD Biosciences), AlexaFluor488 anti-FoxP3 (FJK-16s; Invitrogen), SparkBlue550 anti-CD45 (30-F11; BioLegend), PE anti-RORγt (B2D; Invitrogen), PE/Dazzle594 anti-CD90.2 (30-H12; BioLegend); PerCP-Cy5.5 anti-CD19 (1D3; BD Biosciences), PE-Cy7 anti-IL-17A (eBIO17B7; Invitrogen), BV421 anti-CX3CR1 (SA011F11; BioLegend), eF450 anti-I-A/I-E (M5/114.15.2; Invitrogen), BV570 anti-Ly6C (HK1.4; BioLegend), BV650 andi-CD64 (X54-5/7.1; BD Biosciences), BV711 anti-TNF (XP6-XT22; BD Biosciences), BV750 anti-Ly6G (1A8; BD Biosciences), BV786 anti-SigF (E50-2440; BD Biosciences), BV786 anti-CD3 (17A2; BD Biosciences), BV786 anti-CD19 (1D3; BD Biosciences), FITC anti-CD11b (M1/70; BD Biosciences), PE anti-IL-6 (MP5-20F3; BD Biosciences), PE/Dazzle594 anti-CD103 (2E7; BioLegend), PerCP-Cy5.5 anti-IL-12 (C15.6; BioLegend), PE-Cy7 anti-CD11c (N4/18; BioLegend), BV421 anti-CD8a (53-6.7; BioLegend), BV421 anti-TNF (XP6-XT22; BD Biosciences), AlexaFluor532 anti-CD45 (30-F11; Invitrogen), PE anti-IRF5 (W16007B; BioLegend), PE-Dazzle549 anti-CD64 (X54-5/7.1; BioLegend), PerCP-Cy5.5 anti-SigF (E50-2440; BD Biosciences), APC anti-IL-6 (MP5-20F3; BD Biosciences), AlexaFluor700 anti-Arg1 (A1EXF5; Invitrogen), and APC-Fire750 anti-CX3CR1 (SA011F11; BioLegend). Prior to the acquisition, cells were washed in 1x IMag Stain Buffer (BD Biosciences) and resuspended in stain buffer containing Liquid Counting Beads (BD Biosciences) for cell number calculation. Samples were run in SP6800 Spectral Cell Analyser (Sony Biotechnology, San Jose, CA, USA), in the Nicole Perkins Microbial Communities Core Lab, Snyder Institute for Chronic Diseases, University of Calgary.

### Immunofluorescence staining and analysis

For IF detection and quantification of eosinophils, mLN were excised and fixed overnight in 10% formalin. Paraffin-embedded organs were sectioned by the University of Calgary Diagnostic Services Unit (DSU). IF staining was performed as previously described^130^, with modifications. Slides were deparaffinized and pre-treated at room temperature with Sodium Citrate Buffer (pH 6.0) for 1 hour, Digest-All™ 3 pepsin solution (Thermo-Fisher) for 10 min, and Rodent M Block (BioCare Medical; Pacheco, CA, USA) for 30 min. After three washes, slides were incubated with mouse anti-mouse monoclonal EPX (Clone MM25-8.2.2; Mayo Clinic, AZ, USA) & Rat anti-mouse MBP (Clone MT2.-14.7.3; Mayo Clinic) primary antibody mix at 1:250 overnight at 4°C. After three washes, slides were incubated with goat anti-mouse AF488 (polyclonal; Thermo-Fisher) & goat anti-rat AF647 (polyclonal; Thermo-Fisher) secondary antibody mix at 1:250 at room temperature for 2 hours. After three washes, slides were counterstained with DAPI (Sigma) at 1:1,000 at room temperature for 10 min. The stained slides were imaged in a Nikon A1R laser scanning confocal (Nikon, Tokyo, Japan), in tiles using a 20X 0.75 NA objective. Images were stitched together to visualize the entire section. Eosinophil numbers were quantified using FIJI (ImageJ v.1.54f). Images underwent background subtraction and contrast adjustments, followed by thresholding and subsequent particle analysis using the FIJI BioVoxxel Toolbox plugin. The total tissue area was determined, and the data expressed as eosinophils per mm^2^. Representative data for presentation were imported into Adobe Photoshop v.22.4.2 where brightness and contrast changes were applied to the entire image. All final images were compared to the original raw image to ensure data integrity.

### Data visualization and statistical analysis

Bacterial (shotgun and 16S) and fungal (ITS2) community matrices were merged, and all additional filtering and analysis steps were done using Phyloseq v.1.36.023 ^131^ package. Changes in community structure (beta-diversity) were assessed statistically using PERMANOVA^132^ and visualized in PCoA plots based on Bray-Curtis dissimilarities. Changes in community alpha-diversity and species richness were measured using the Shannon and Chao1 indexes, respectively. To further visualize the co-occurring changes in community diversity in ANTIBIO participants, normalized scaled observed fungal and bacterial alpha-diversity were plotted per individual participant. To explore the changes in taxonomic community structure, boxplots were made to identify outliers and normality was tested using the Shapiro-Wilk test. If data were normally distributed (*P* > 0.05), significant changes in relative abundance were determined with unpaired t-test, and if data were not normally distributed (*P* ≤ 0.05), significant changes were determined with Mann-Whitney (Wilcoxon rank sum) test. The delta relative abundance for the most abundant ASVs was determined as the change between relative abundance values per post- and pre-antibiotic samples. DESeq2 v.1.38.3 ^133^ was used to determine differentially abundant ASVs in the shotgun and ITS2 dataset between time points. We used the NetCoMi R package v.1.1.0 ^50^ to build interkingdom microbial association networks using the ITS2 and shotgun datasets. Bacterial and fungal co-occurrence networks were constructed using relative abundance with variance stabilizing transformation at the species level, a Pearson correlation threshold of 0.5, and a fast greedy clustering algorithm. Hub taxa were identified as those having the highest betweenness centrality, associated with high connectivity with different nodes and sub-networks^50^. Pair-wise network comparisons were made between pre- and post-antibiotic time points for measures of centrality and hub taxa, calculated based on 5,000 permutations. Random forest was performed to determine the most relevant factors (infant, pregnancy and delivery, and antibiotic treatment) predictive of *Malassezia* spp. Expansion. Multivariable logistic regression analysis was performed using package lme4 v.1.1.33 ^134^ to identify factors associated with an increase in *Malassezia* abundance. Only the top five predictors identified by random forest were included in the logistic regression analysis to prevent model overfitting. Results are reported as log-transformed OR and 95% CI for each factor. The relative abundance values for the *Malassezia* genus on day 0 and follow-up samples were used to define a *Malassezia* trend variable (“expansion”: *n*=23, “no-expansion”: *n*=13) for each participant. The “no-expansion” variable was set as a reference level for both random forest and logistic regression analyses. Fecal metabolite changes between colonization groups were visualized in a PCA plot and a dendrogram of the top 50 most abundant metabolites. To determine the significant differences in metabolite detection, we used volcano plots, combining results from FC analysis and t-test, with an FC and *P*-value threshold set to 2 and 0.05, respectively. PCA, dendrogram, t-tests, and volcano plot analysis of fecal metabolites from gnotobiotic mice were performed using MetaboAnalyst v.5.0 ^126^. Flow cytometry data obtained in the SP6800 Spectral Cell Analyser (Sony) were exported as .fcs files and analyzed using FlowJo v.10.8.1 (https://flowjo.com). Graphs were made and analyzed using Prism v.10.0.2 (GraphPad Software Inc.). Bar graphs display the mean with standard error of the mean (S.E.M.), and all data points represent biological replicates. Outliers were identified using the ROUT test (Q=1%) and removed. Statistically significant differences between colonization groups were determined with t-test or ANOVA. For all tests, significance was set at *P*<0.05.

## List of Supplementary Materials

Materials and Methods

Fig. S1 to S10

Table S1 to S7

## Acknowledgments

We thank all ANTIBIO participants and families that took part in this study. Dr. Elizabeth Jacobsen for kindly providing the anti-mouse EPX and MBP antibodies. Dr. Bruna Araujo and Dr. Matheus Carneiro for assistance with the flow gating strategy. Dr. Aline Ignacio, Dr. Lukas Mager, and Dr. Fernanda Castanheira for gut, mLN, and lung digestion protocol support. Jared Schlechte and Dr. Michelle Asbury for help with microbiome analysis. Dr. Mona Parizadeh with support depositing the shotgun files to NCBI. Dr. Margaret Kelly and Elaine de Heuvel for histopathology and lung scoring training. Panels in figures 1, 3, 4, S7-10 created with biorender.com.

## Funding

EvTB was funded by the Eye’s High Doctoral Recruitment Scholarship and the Canadian Institutes for Health Research (CIHR) Frederick Banting and Charles Best Canada Graduate Scholarships Doctoral Award. MCA is supported by funds from the Cumming School of Medicine, the Alberta Children Hospital Research Institute, the Snyder Institute of Chronic Diseases, and the CIHR (Grant 180429).

## Author contributions

Conceptualization: EvTB, SBF, MCA

Study Design: EvTB, MWG, WNTN, KDP, KDM, SBF, MCA

Clinical data processing and analysis: EvTB, MWG, EM, HRR

Mouse experiments: EvTB, MWG, WNTN, EMM, TG, KK

Sample processing and data analysis: EvTB, MWG, WNTN, EMM, TG, CAT, NG, KK

Microbiome data processing and analysis: EvTB, MG, EMM, HRR, TH

Immunofluorescence staining and analysis: WNTN

Funding acquisition: SBF, MCA

Writing – original draft: EvTB, MCA

Writing – review & editing: EvTB, MWG, WNTN, EMM, HRR, TG, CAT, TH, NG, KK, KDP, KDM, SBF, MCA

## Competing interests

Authors declare that they have no competing interests.

## Data and materials availability

Demultiplexed shotgun (ANTIBIO), ITS2 (ANTIBIO), and 16S (gnotobiotic mice) sequencing files were deposited into the Sequence Read Archive (SRA) of NCBI and can be accessed via BioProject accession numbers PRJNA1055141, PRJNA1053397, and PRJNA1053409, respectively. All codes (R scripts) have been deposited at: https://github.com/ArrietaLab. Additional sequencing and immunological data are provided in a Supplementary Source Data File. All microbial strains used are kept as stocks in the Arrieta lab and can be shared when requested. Any additional information required to reanalyze the data reported in this paper is available from the lead contact upon request.

## Notes

### Competing Interest Statement

The authors have declared no competing interest.

