## Supplemental Material for "Antibiotic-induced *Malassezia* spp. expansion in infants promotes early-life immune dysregulation and airway inflammation in mice"

van Tilburg Bernardes *et al.*

Corresponding author: Dr. Marie-Claire Arrieta,

**This PDF file includes:**

Supplemental Materials and Methods

Fig. S1 to S10

Tables S1 to S6

**Other Supplementary Material for this manuscript includes the following:**

Microbiome analysis codes (lab Github)

ANTIBIO study participant metadata (source data file)

Murine model data (source data file)

**SUPPLEMENTAL MATERIALS AND METHODS:**

**ANTIBIO study design**

*Sample size calculation*

Based on data from the infant study performed in Ecuador ^1^, the required sample size to detect significant fungal changes in association with antibiotic use was determined using MetSizeR ^2^. MetSizeR is a publicly available software package in R designed to estimate sample size for high-throughput experiments. For the sample size calculation, we assumed at least 100 parameters were measured and that 10% would differ significantly between short- and long-term groups. This approach demonstrated that a sample size of 44 was required to maintain a false discovery rate below 0.05. Therefore, we designed this study to enroll approximately 45 participants.

*Patient screening*

Potential participants were identified in the ACH-ED by a Research Assistant from the Pediatric Emergency Medicine Research Assistant Program (PEMRAP) or by a Research Nurse from the PERT. The Research Assistant or Research Nurse scanned SEC (tracking program for patients in the ED) for patients who may qualify for the study. All patients presenting to the ED are entered into this system at triage. At the initial screening, the study coordinator explained the nature of the research project to the infant’s caregivers. After informed consent was obtained, a baseline assessment was documented to ensure the subject’s eligibility for inclusion in the study.

*Inclusion criteria*

To meet eligibility, patients needed to be: (i) admitted as an in-patient to the ACH, the Peter Lougheed Centre, or sent to the Home or Ambulatory Parental Therapy Program; (ii) administered an intravenous (IV) dose of antibiotics; and (iii) younger than 6 months of age (0 - 5.99 months).

*Exclusion criteria*

Subjects were excluded if they: (i) were currently taking oral antibiotics or have been taking oral antibiotics within the past 2 weeks; (ii) were born very prematurely (before 32 weeks of gestation); (iii) had any congenital gastrointestinal anomalies or had a history of gastrointestinal surgery; (iv) had a history of necrotizing enterocolitis; (v) had a history of cholestasis (i.e.: liver dysfunction) or currently experiencing gastroenteritis (i.e.: diarrhea); or (vi) are currently known to be neutropenic (absolute count < 1,000 neutrophils/mm^3^), or at high-risk of being neutropenic (i.e.: receiving chemotherapy). Besides the aforementioned exclusion criteria, any infants (vii) residing outside of the Calgary Metropolitan Zone, (viii) unable to provide all requested samples, (ix) who failed to complete follow-up, or (x) were unable to understand English were also excluded from the study.

*Patient enrolment and follow-ups*

Following initial screening and confirmation of all inclusion criteria, the study coordinator explained the nature of the research project to the infant’s caregiver. Informed consent had to be obtained prior to the child being enrolled in the study. A detailed medical history was documented to ensure the subject’s eligibility for inclusion in the study. Descriptive data regarding previous antibiotic use, feeding practices (i.e.: breast milk, formula, or both), household pets, number of siblings, previous infections, etc. were collected in the enrolment questionnaires. Medical records were reviewed to obtain necessary data. Patients’ charts were also reviewed in the ED. All information was entered into the University of Calgary’s Research Electronic Data Capture (REDCap) system.

Caregivers were contacted on day 2 post-enrolment and asked about the duration of antibiotic treatment to coordinate final follow-up. When necessary, we contacted the caregivers again until the precise duration of antibiotic treatment was known. If the anticipated duration of antibiotic therapy was ≥ 2 days, caregivers were reminded to collect a specimen (fecal sample or rectal swab) within 24 hours of the last dose of antibiotic(s) using supplies in a sample collection kit provided before discharge. On the last day of antibiotic treatment, which was anytime between day 2 and day 14, the participant’s caregiver was reminded to collect the fecal sample. Also, at follow-up, clinical information about ongoing symptoms or complications (i.e.: fever, diarrhea, diaper rash, and oral thrush) was obtained from the caregiver and recorded into REDCap.

*Sample storage and delivery*

For samples collected in the ACH-ED, nurses immediately delivered them to the Alberta Precision Laboratory (APL) at ACH, a subsidiary of Alberta Health Services, for immediate freezing at -80^o^C. Samples were then retrieved by a lab member for transport and continuous storage at -80^o^C until further processing. For samples collected at home, the subject’s caregivers were instructed to place the fecal samples in a -20^o^C freezer until transport to the lab. Caregivers were given the contact for a courier service that retrieved the sample from their house to bring it directly to the Arrieta lab, where it was received and immediately stored at -80^o^C. This transport chain has been previously performed in the Alberta Provincial Pediatric Enteric Infection Team (APPETITE) study (REF).

**Microbial isolation from ANTIBIO fecal samples and isolate characterization**

Samples from patients that demonstrated fungal and/or *Malassezia* spp. overgrowth were selected for fungal isolation. Approximately 100 mg of frozen fecal samples were dissolved in 500 μL of sterile saline (0.9% NaCl), serially diluted, and plated in YM (BD) or modified Dixon (mDixon) ^3^ agar plates containing chloramphenicol (200 mg/L; Sigma) and gentamicin (10 mg/L; Sigma). Plates were incubated at room temperature (~25°C) and monitored for growth for up to 10 days. Morphologically distinct colonies grown in a plate were retrieved and characterized with colony PCR, as previously described ^4-6^. The 16S gene and the ITS region were selected for bacterial and fungal characterization, respectively. 16S gene was amplified using 8FW (5’-AGAGTTTGATCCTGGCTCAG-3’) and 926RV (5’- CCGTCAATTCCTTTRAGTTT -3’) primers. The ITS region was amplified using ITS-FW (5′- GTGAATCATCGAATCTTTGAA -3′) and ITS-RV (R-5′-TCCTCCGCTTATTGATATGA-3′) primers. 16S and ITS PCR were run in 25 μL reactions, containing 12.5 μL TopTaq Master Mix kit (Qiagen), 1 μL of each primer at 10 μM, 1 μL of boiled colonies supernatant, and 9.5 μL of nuclease-free water. PCR thermocycler program consisted of an initial 5 min denaturation step at 95°C, followed by 35 cycles of 95°C for 30 s, 55°C (16S) or 50°C (ITS) for 30 s, and 72^o^C for 60 s, and a final step of 72^o^C for 7 min. PCR products were visualized in 1.5% agarose gel, and cleaned up using the QIAquick PCR purification kit (Qiagen), following manufacturer’s instructions. Sanger sequencing was performed at the Center for Health Genomics and Informatics, at the University of Calgary. Returned sequences were used for the Nucleotide Basic Local Alignment Search Tool (BLASTN) similarity search (<https://www.ncbi.nlm.nih.gov/BLAST>). Identification was determined when a percent sequence identity of > 95% was achieved.

**16S qPCR**

The 16S qPCR runs were performed as previously described ^7^. Briefly, 10-μL reactions consisted of 5 μL of 2x iQ SYBR Green Supermix (BioRad Laboratories), 0.5 μL of u16Sfr primer (5’-TCC TAC GGG AGG CAG CAG T-3’) at 10 μM, 0.5 μL of u16Srv primer (5’- GGA CTA CCA GGG TAT CTA ATC CTG TT-3’) at 10 μM, 2 μL of nuclease-free water, and 2 μL of 1 ng/μL dilution of extracted template DNA. The 16S qPCR thermocycler program consisted of an initial 5 min step at 94^o^C, followed by 40 cycles of 94^o^C for 15 s, 60^o^C for 30 s, and 72^o^C for 30 s, and a final melt curve consisting of a single cycle of 95^o^C for 15 s, 60^o^C for 1 min, 95^o^C for 15 s, and 60^o^C for 15 s.

**ITS qPCR**

The ITS qPCR runs were performed as previously described ^8^, with modifications. For fungal biomass quantification in infant fecal samples, ITS qPCR 20-μL reactions consisted of 10 μL of 2x iQ SYBR Green Supermix (BioRad Laboratories), 2 μL of ITS1F primer (5’- CTTGGTCATTTAGAGGAAGTAA-3’) at 1 μM, 1 μL of ITS4 primer (5’- TCCTCCGCTTATTGATATGC-3’) at 10 μM, 5 μL of nuclease-free water, and 2 μL of 1 ng/μL dilution of extracted template DNA. The ITS qPCR thermocycler program consisted of an initial 5 min step at 95 °C, followed by 40 cycles of 95°C for 30 s, 55°C for 30 s, and 72°C for 1 min, and a final melt curve consisting of a single cycle of 95°C for 15 s, 60°C for 1 min, 95°C for 15 s, and 60°C for 15 s. For *M. restricta* quantification in murine whole colon contents, to account for the lower fungal abundance, the ITS qPCR 20-μL reactions were modified to include 2 μL of 10 ng/μL dilution of extracted template DNA and the thermocycler run was changed to include 40 amplification cycles.

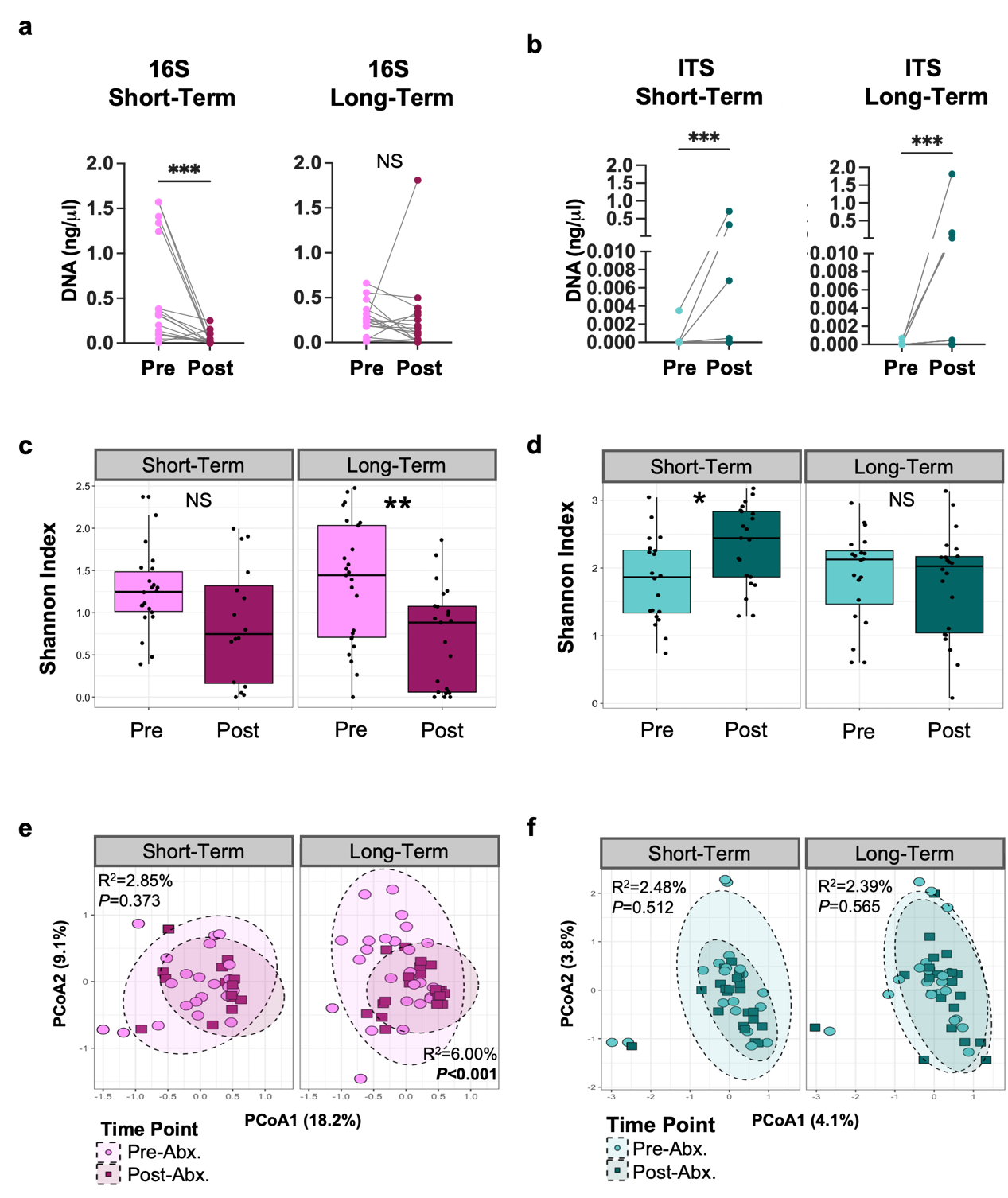

**Fig. S1: Antibiotic impact on infant microbiome biomass and diversity: (a)** 16S qPCR quantification (standard curve method) of bacterial biomass in ANTIBIO samples from pre- and post-antibiotic treatment from the short and long-term samples. **(b)** ITS qPCR quantification of fungal biomass in short and long-term samples. **(c)** Shannon index plots of bacterial alpha diversity. **(d)** Shannon index plots of fungal alpha diversity. **(e)** PCoA plots of bacterial beta-diversity on Bray-Curtis dissimilarities across time points and treatment groups. **(f)** PCoA plots of fungal beta-diversity on Bray-Curtis dissimilarities across time points and treatment groups. (c-d) Boxplots indicate median (inside line) and 25th and 75th percentile as the lower and upper hinges, respectively. (a, b, c, e) Statistically significant differences defined by t-test or paired Wilcoxon test. **P*<0.05, ***P<*0.01, ****P<*0.001, NS=not significant. (e-f) Ellipses represent a 95% confidence interval from data points. Statistical analysis performed with PERMANOVA.

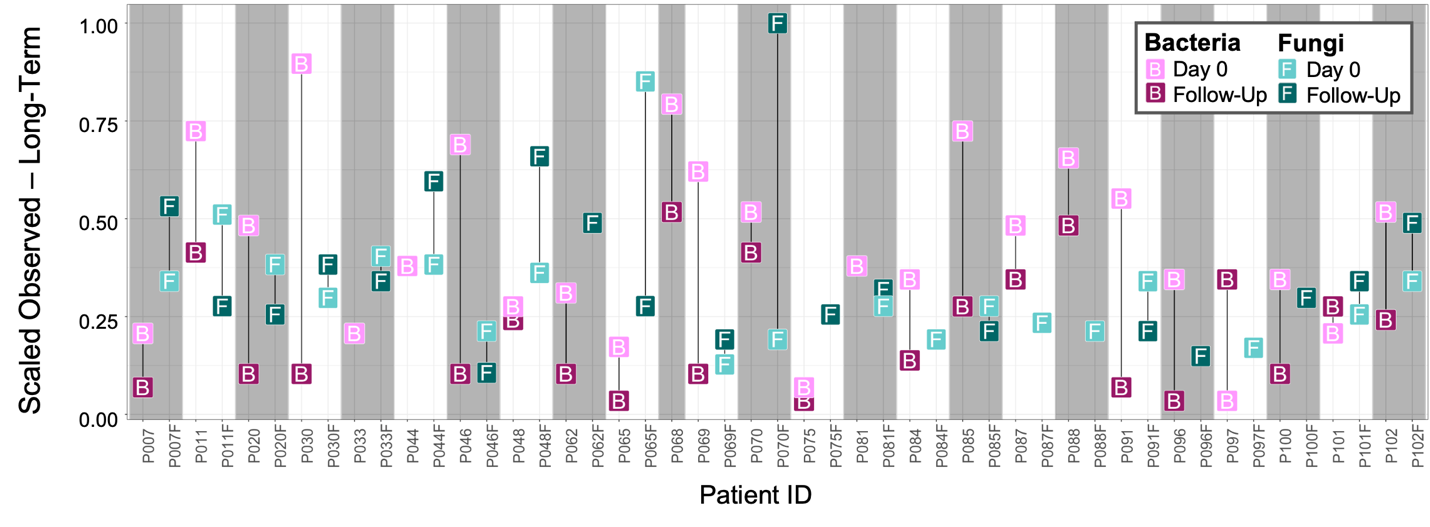

**Fig. S2: Shifts in fungal and bacterial alpha diversity per participant receiving long-term antibiotic.** Scaled observed diversity for fungal and bacterial assignments in the long-term treatment group per participant. Values for each participant enrolled in the study are indicated by alternated shading color.

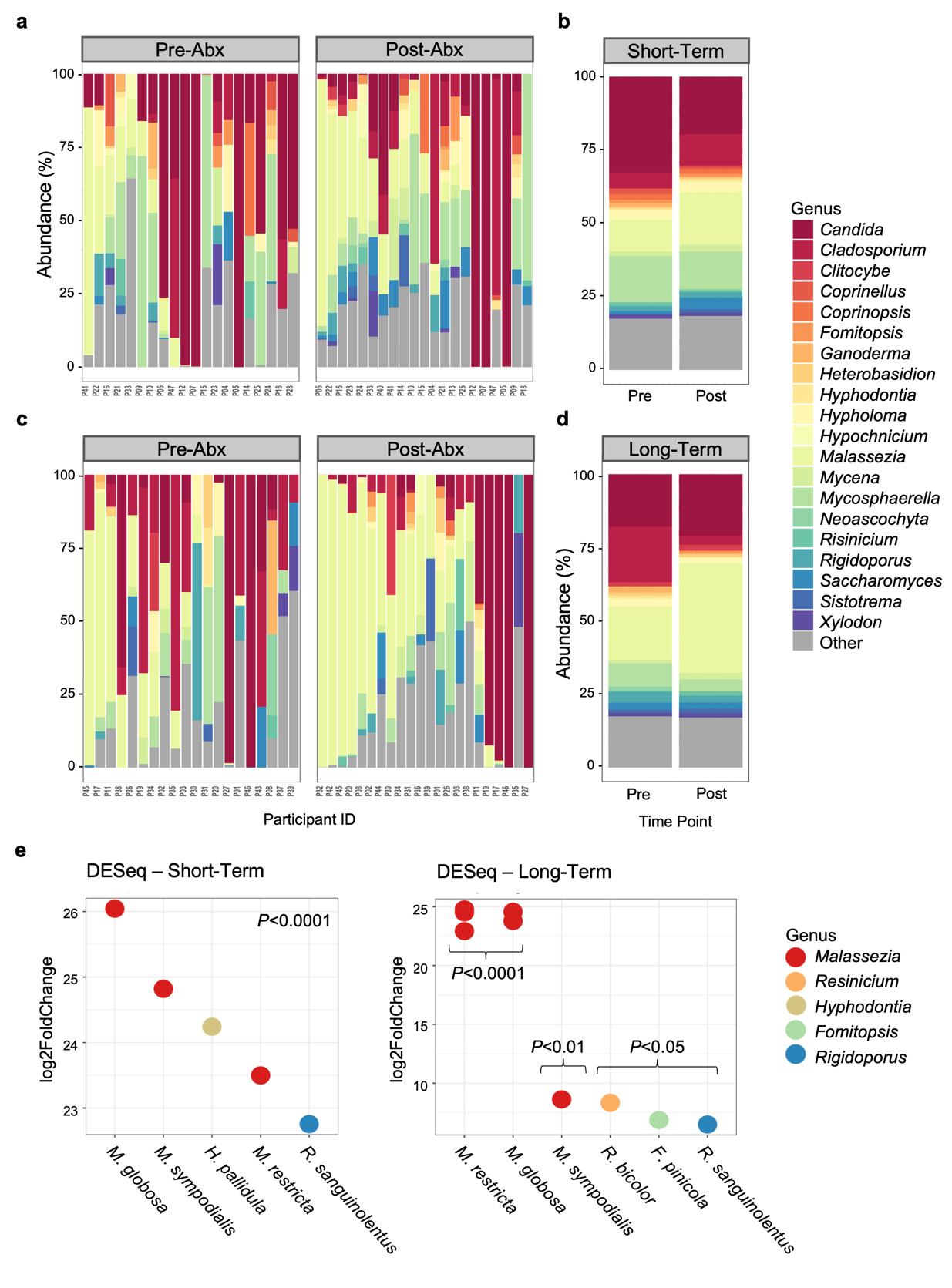

**Fig. S3: Antibiotic impact on infant mycobiome composition. (a)** Individual and **(b)** median relative abundance for the 20 most abundant fungal taxa for participants in the short-term group. **(c)** Individual and **(d)** median relative abundance for the 20 most abundant fungal taxa for participants in the long-term group. (a-d) Taxa merged at the genus level. **(e)** Differentially detected fungal ASV in the short- and long-term groups. (e) Statistically significant changes in ASV detection between pre- and post-antibiotic determined by DESeq2 and FDR corrected; *Padj*<0.05.

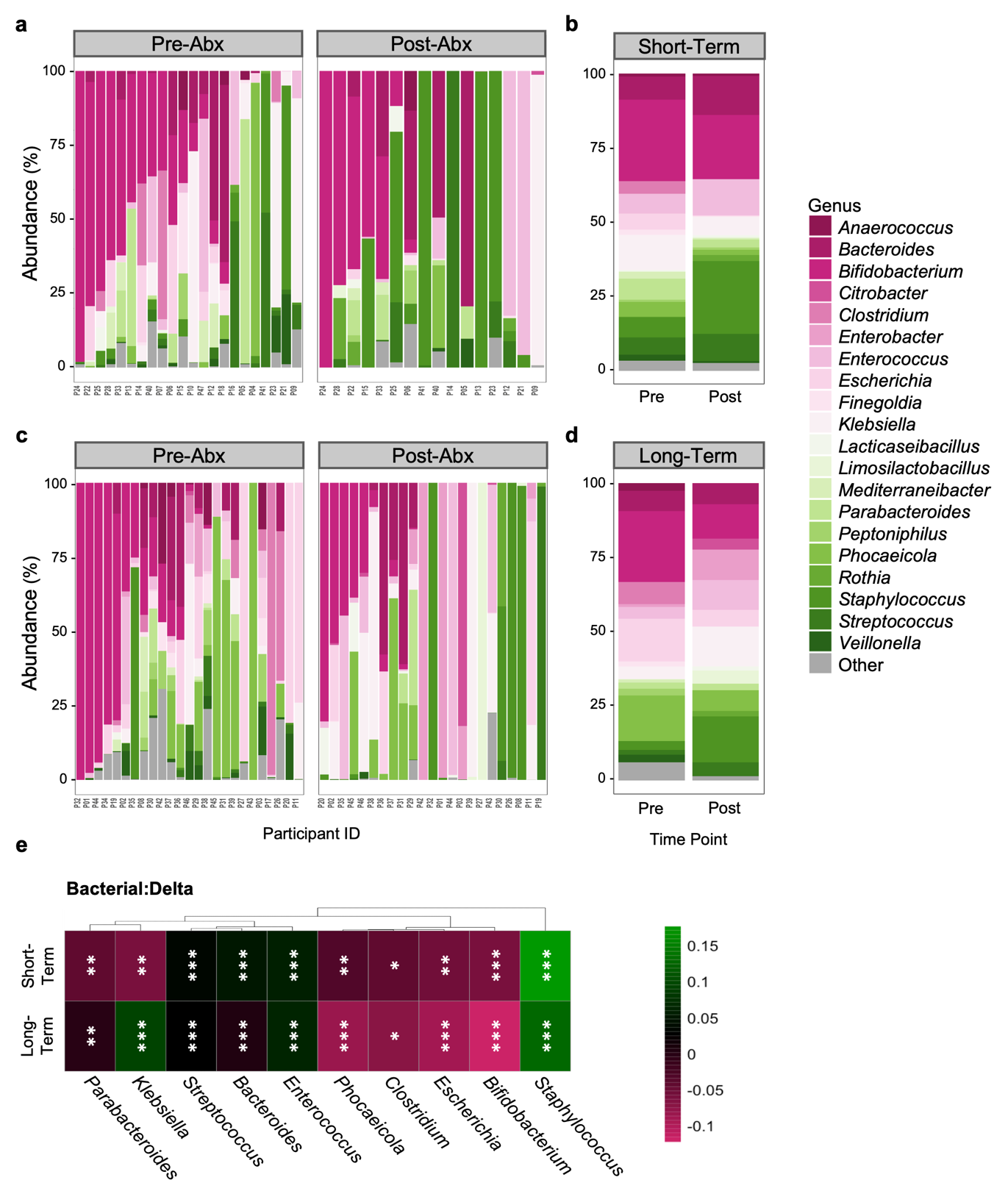

**Fig. S4: Antibiotic impact on infant microbiome composition: (a)** Individual and **(b)** median relative abundance for the 20 most abundant bacterial taxa for participants in the short-term group. **(c)** Individual and **(d)** median relative abundance for the 20 most abundant bacterial taxa for participants in the long-term group. (a-d) Taxa merged at the genus level. **(e)** Delta relative abundance changes in the top 10 most abundant bacterial genera identified in the shotgun dataset per study arm. Color denotes increase (green) and decrease (pink) abundance following antibiotic treatment (pre- vs*.* post-antibiotic). Statistically significant differences defined by paired sample Wilcoxon test: **P*<0.05, ***P<*0.01, ****P<*0.001.

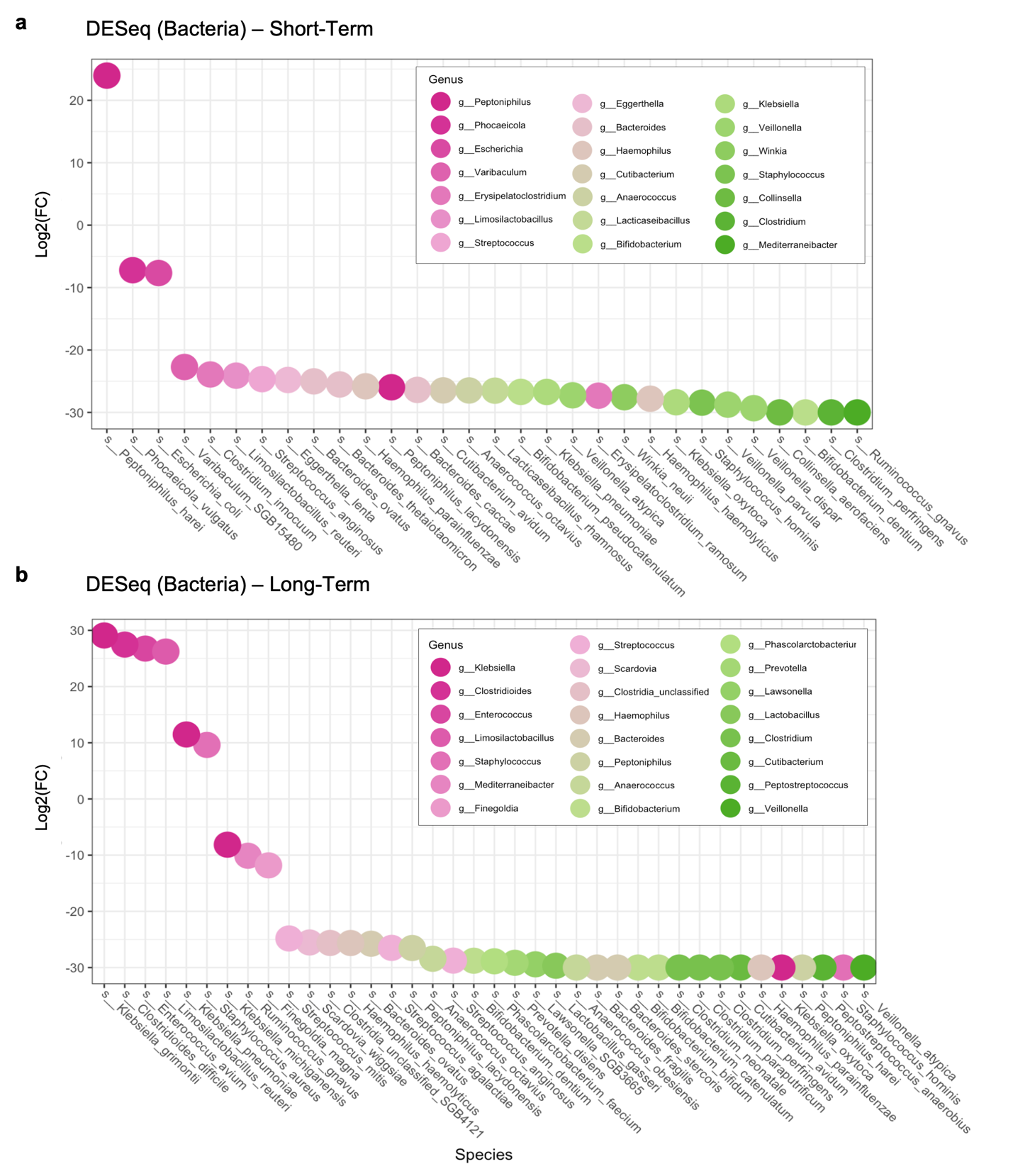

**Fig. S5: Differentially detected bacterial species in antibiotic-treated samples.** Significant log2 FC detected in bacterial abundance in the **(a)** short- and **(b)** long-term groups. Statistically significant changes in species detection between pre- and post-antibiotic treatment determined by DESeq2 and FDR corrected; *Padj*<0.05.

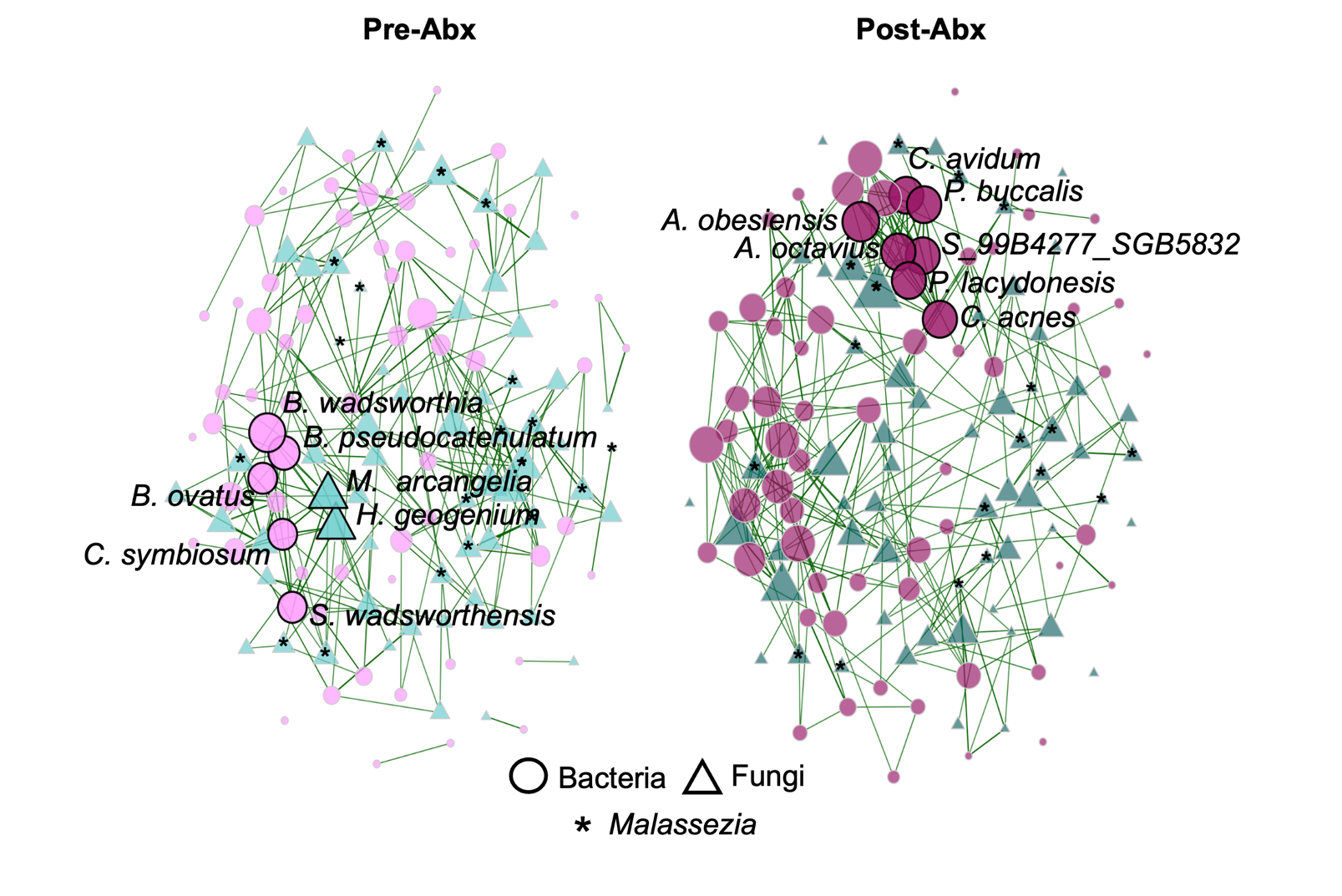

**Fig. S6: Estimated fungal-bacterial correlation networks in infant microbiome.** Comparison of bacterial and fungal correlation networks in pre- (*N*=40) and post-antibiotic treatment (*N*=35) samples. Samples from either treatment group (i.e.: short- or long-term), were combined to determine the impact of antibiotic treatment on microbiome associations. Bacterial and fungal inter-kingdom co-occurrence network analysis was performed at the ASV level (ITS) and bacterial assignment (shotgun). Pair-wise network comparisons were made between pre- and post-antibiotic time points using relative abundance with variance stabilizing transformation. A Pearson correlation coefficient threshold of 0.5 was applied. Bacteria and fungi are defined by circle and triangle shapes, respectively. Members of the *Malassezia* genus within networks are marked with an asterisk. Hubs are highlighted by bold text and borders. Estimated correlations drawn with green edges. The layout was kept the same for both networks.

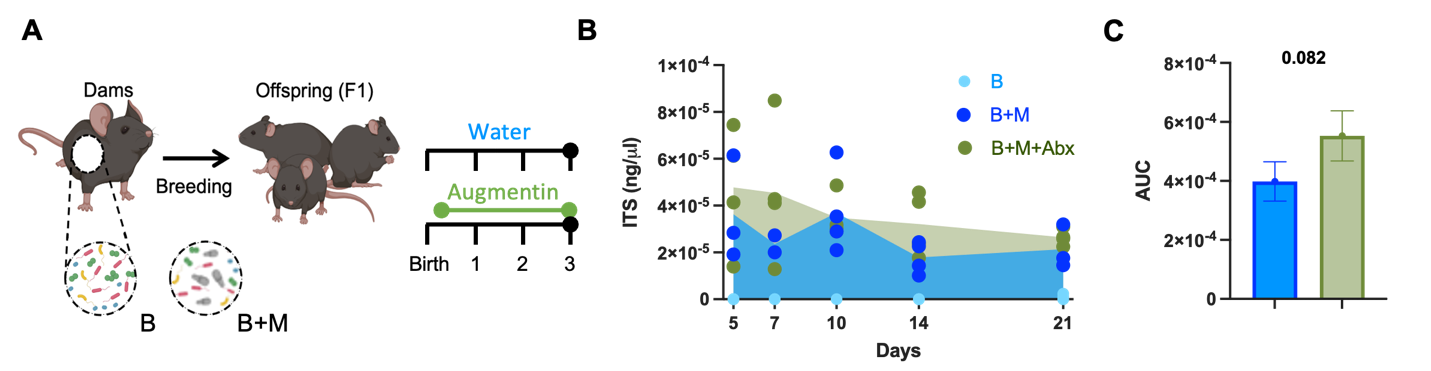

**Fig. S7: *M. restricta* DNA quantification in gnotobiotic mice.** *M. restricta* colonization was determined in gnotobiotic mice throughout neonatal development. **(a)** Experimental layout for early-life *M. restricta* colonization model. Germ-free dams received two gavages with colonization consortia containing 12-mouse derived bacteria (B), or bacteria + *M. restricta* (B+M). A group of B+M mice was further treated with the antibiotic Augmentin (B+M+Abx) from the days 2 to 21 post-birth. **(b)** ITS qPCR quantification (standard curve method) of *M. restricta* DNA in whole colon contents from day 5 to 21 post birth. **(c)** Cumulative *M. restricta* quantification with area under the curve (AUC) analysis. Data represented as the mean of total peak area ± S.E.M. (b-c) Color denotes colonization treatment (B = light blue, B+M = royal blue, B+M+Abx = green). All data points represent biological replicates *(N=*3-4/group/time point). (c) Statistical analysis was performed with T-test.

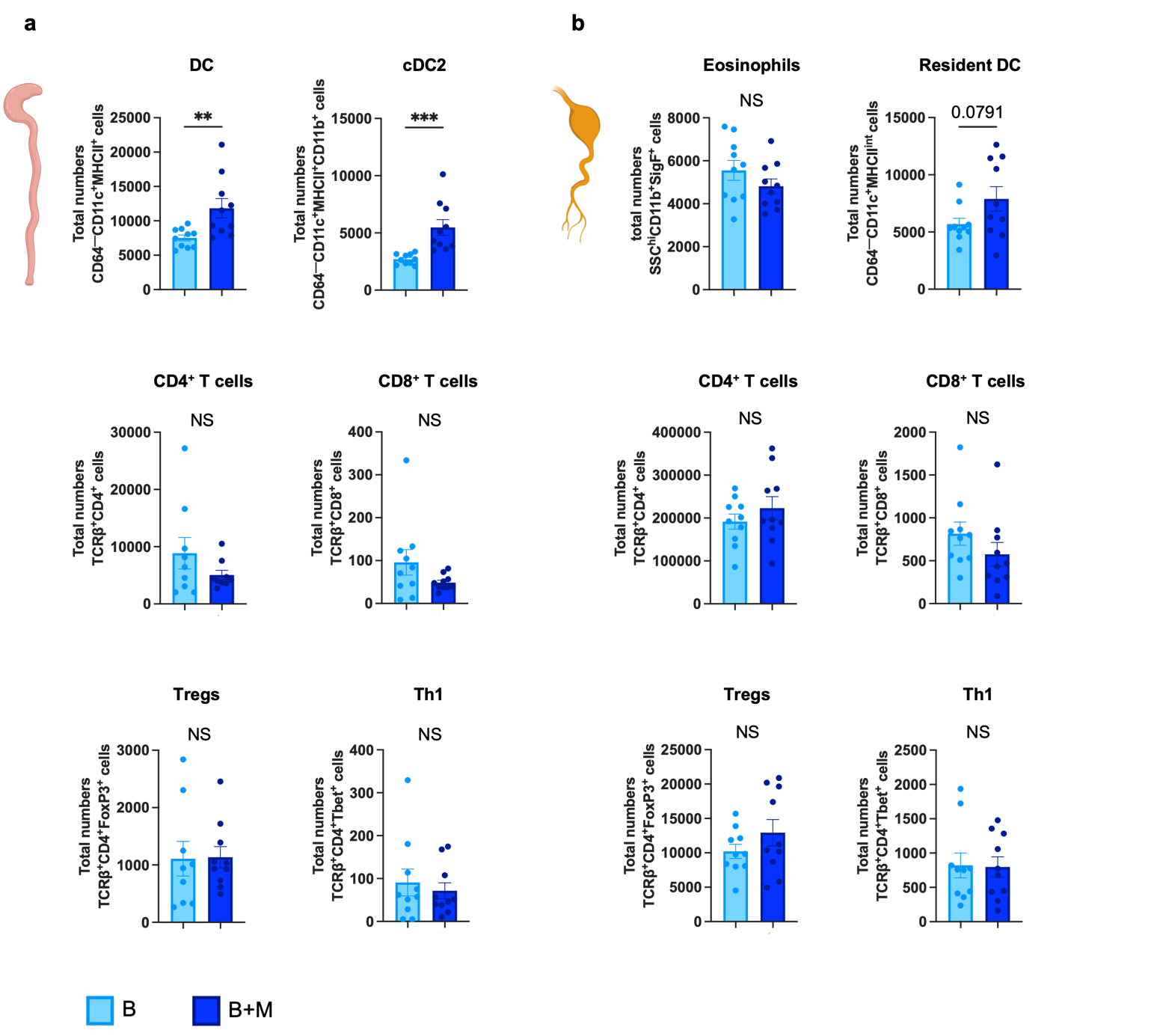

**Fig. S8: Additional early-life immune readouts in *M. restricta* colonized gnotobiotic mice.** Immune functions in cLP and mLN were determined at 3 weeks of age in F1 mice. **(a)** Total numbers of DC, cDC2, CD4^+^ T cells, CD8+ T cells, Tregs, and Th1 cells in cLP. **(b)** Total numbers of eosinophils, resident DC, CD4^+^ T cells, CD8+ T cells, Tregs, and Th1 cells in mLN. Data represented as mean ± S.E.M. All data points represent biological replicates (cLP, mLN, *N*_B_=10, *N*_B+M_=10). Color denotes colonization treatment (B = light blue, B+M = royal blue). Statistically significant differences defined by T-test; ***P* <0.01, ****P* <0.001, NS = not significant.

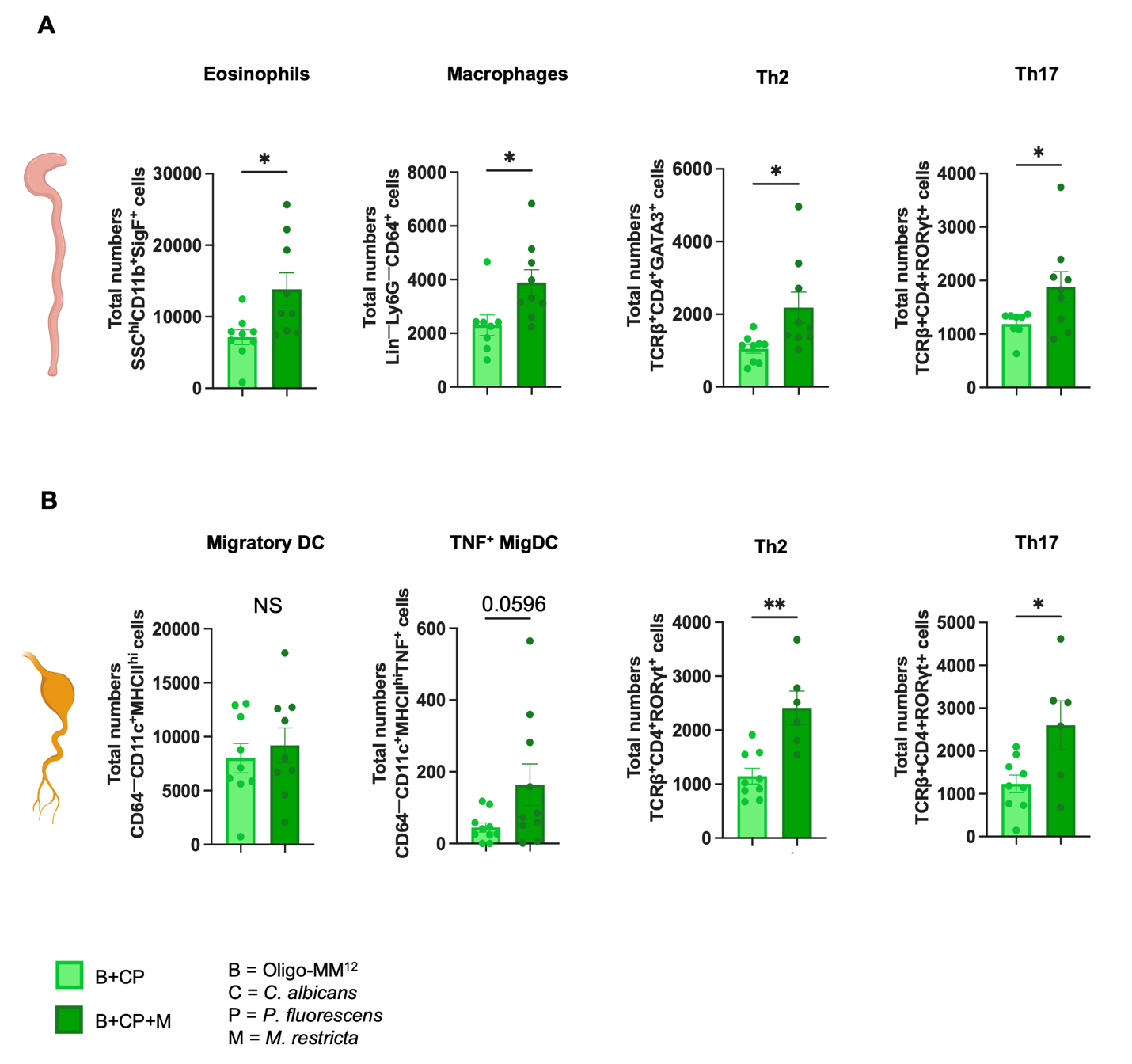

**Fig. S9: Early-life *M. restricta* exposure induces immune changes in gnotobiotic mice containing other immunogenic colonizers.** Gnotobiotic mice were colonized with Oligo-MM12, *C. albicans* (isolate K3), and *P. fluorescens* (J4) (B+CP) or also with *M. restricta* (B+CP+M) from birth. Immune functions in cLP and mLN were determined at 3 weeks of age. **(a)** From left to right, total cell numbers of eosinophils, macrophages, Th2, and Th17 in cLP. **(b)** From left to right, total cell numbers of migratory DC (migDC), TNF^+^ MigDC, Th2, and Th17 in mLN. Data represented as mean ± S.E.M. All data points represent biological replicates (*N*_B+CP_=9, *N*_B+CP+M_=9). Color denotes colonization treatment (B+CP=light green, B+CP+M=dark green). Statistically significant differences determined by T-test; **P*<0.05, ***P*<0.01, NS=not significant.

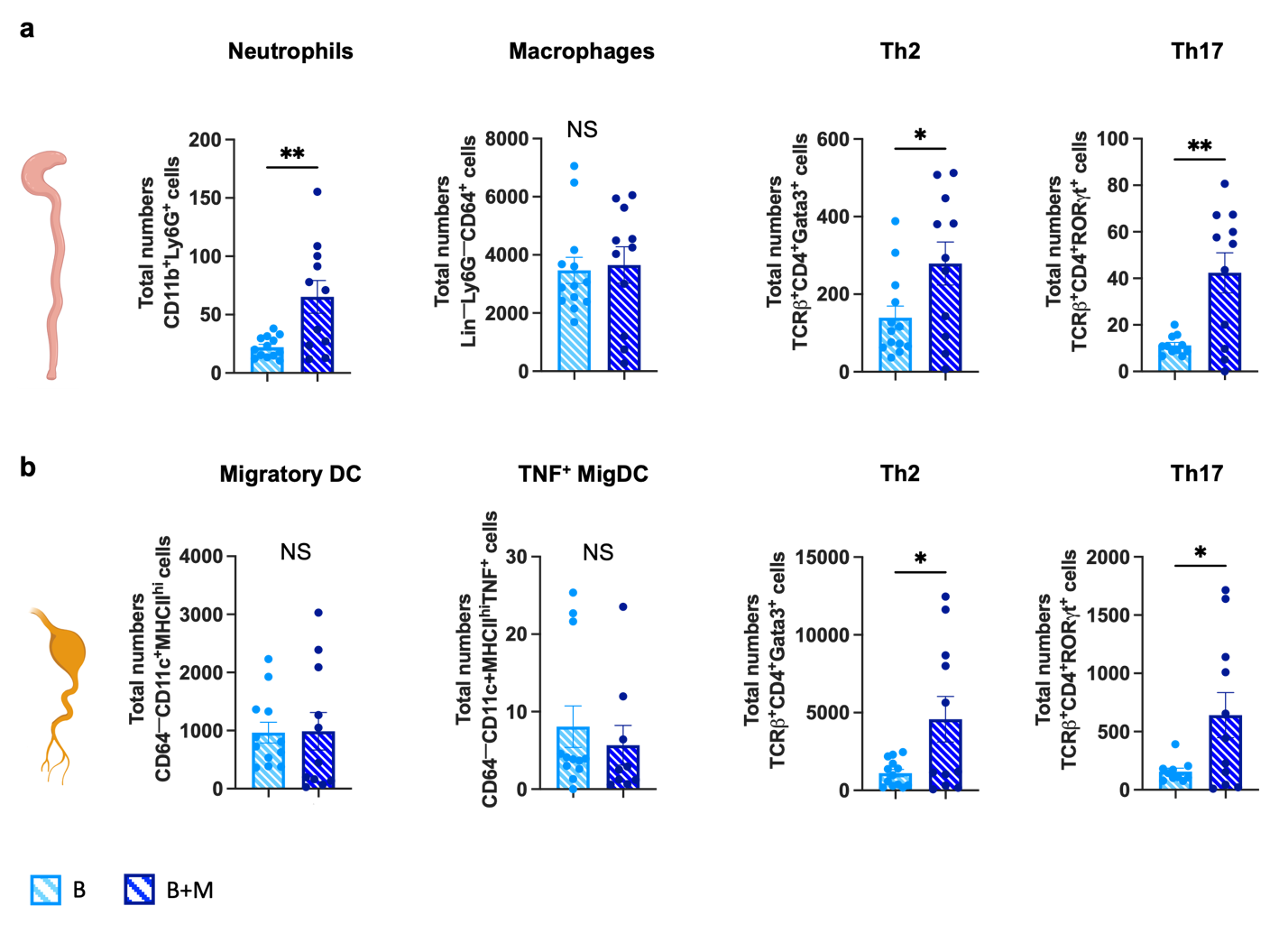

**Fig. S10: *M. restricta*-induced intestinal immune changes in *ΔdblGATA1* mice.** Immune functions were determined in cLP and mLN of *ΔdblGATA1* gnotobiotic mice at 3 weeks of age. **(a)** From left to right, total cell numbers of neutrophils, CD64^+^ macrophages, Th2, and Th17 in cLP. **(b)** From left to right, total cell numbers of migDC, TNF^+^ MigDC, Th2, and Th17 in mLN. Data represented as mean ± S.E.M. All data points represent biological replicates (*N*_B_=13, *N*_B+M_=11). Color denotes colonization treatment (B=streaked out light blue, B+M=streaked out royal blue). Statistically significant differences determined by T-test; **P*<0.05, ***P*<0.01, NS=not significant.

**Table S1: ANTIBIO study participant characteristics.**

| **Study Variables** | | | **Short-Term**  **(n = 22)** | **Long-Term**  **(n = 25)** | ***P-*value**^Φ^ |
| --- | --- | --- | --- | --- | --- |
| **Child** | | |  |  |  |
|  | Sex (Female) | | 6 (27.3%) | 10 (40.0%) | 0.3582 |
|  | Median age in days (range) | | 21.5 (3-108) | 33.0 (3-133) | 0.1574 |
|  | Median weight at birth in Kg (range) | | 3.4 (2.35-5.60) | 2.8 (1.87-3.88) | **0.0217** |
|  | Breast milk within 1h after delivery (Yes) | | 14 (63.6%) | 18 (72.0%) | 0.5394 |
|  | Formula within first week after delivery (Yes) | | 12 (54.5%) | 20 (80.0%) | 0.0933 |
|  | Current feeding | |  |  |  |
|  |  | Breast milk | 10 (45.5%) | 12 (48.0%) | 0.6005 |
|  |  | Formula | 5 (22.7%) | 8 (32.0%) |  |
|  |  | Both | 7 (31.8%) | 5 (20.0%) |  |
|  | Antibiotics since birth (Yes) | | 2 (9.1%) | 2 (8.0%) | 0.8936 |
|  | Previous infections | |  |  |  |
|  |  | Gastrointestinal | 1 (4.5%) | 0 (0.0%) | 0.2812 |
|  |  | Ear | 1 (4.5%) | 0 (0.0%) | 0.2812 |
|  |  | Urinary | 0 (0.0%) | 1 (4.0%) | 0.3636 |
| **Pregnancy and delivery** | | |  |  |  |
|  | Pregnancy complications | |  |  |  |
|  |  | Diabetes | 2 (9.1%) | 4 (16.0%) | 0.4788 |
|  |  | Hypertension | 0 (0.0%) | 1 (4.0%) | 0.3430 |
|  |  | Preeclampsia | 1 (4.5%) | 0 (0.0%) | 0.2812 |
|  |  | Others | 6 (27.3%) | 5 (20.0%) | 0.5568 |
|  | Antibiotics during pregnancy (Yes) | | 1 (4.5%) | 2 (8.0%) | 0.6032 |
|  | Type of delivery | |  |  |  |
|  |  | Caesarian | 6 (27.3%) | 6 (24.0%) | 0.7974 |
|  |  | Caesarian planned | 2 (9.1%) | 2 (8.0%) | 0.9999 |
|  |  | Median gestational age in weeks (range) | 38.5 (35-41) | 38.0 (33-40) | 0.3747 |
|  | Antibiotics during labour (Yes) | | 4 (18.2%) | 4 (16.0%) | 0.8923 |
|  | Group B Strep status at delivery | |  |  |  |
|  |  | Positive | 3 (13.6%) | 3 (12.0%) | 0.9455 |
|  |  | Unknown | 3 (13.6%) | 5 (20.0%) | 0.5624 |
| **Household** | | |  |  |  |
|  | Pets | |  |  |  |
|  |  | No | 8 (36.4%) | 17 (68.0%) | **0.0301** |
|  |  | Cat(s) | 2 (9.1%) | 1 (4.0%) | 0.4762 |
|  |  | Dog(s) | 10 (45.5%) | 6 (24.0%) | 0.1214 |
|  |  | Cat(s) and dog(s) | 2 (9.1%) | 1 (4.0%) | 0.4762 |
|  |  | Others | 2 (9.1%) | 0 (0.0%) | 0.1234 |

| **Study Variables (Continued)** | | | **Short-Term**  **(n = 22)** | **Long-Term**  **(n = 25)** | ***P-*value**^Φ^ |
| --- | --- | --- | --- | --- | --- |
| **Household** | | |  |  |  |
|  | Siblings | |  |  |  |
|  |  | 0 | 8 (36.4%) | 8 (32.0%) | 0.9118 |
|  |  | 1 | 8 (36.4%) | 14 (56.0%) | 0.1782 |
|  |  | 2 | 4 (18.2%) | 2 (8.0%) | 0.2966 |
|  |  | ≥3 | 2 (9.1%) | 1 (4.0%) | 0.4762 |
| **Chart review from triage** | | |  |  |  |
|  | CTAS score | |  |  |  |
|  |  | 1 | 2 (9.1%) | 4 (16.0%) | 0.4788 |
|  |  | 2 | 17 (77.3%) | 14 (56.0%) | 0.1246 |
|  |  | 3 | 2 (9.1%) | 6 (24.0%) | 0.1748 |
|  |  | 4 | 1 (4.5%) | 0 (0.0%) | 0.2812 |
|  |  | 5 | 0 (0.0%) | 1 (4.0%) | 0.3430 |
|  | Mean first temperature at ED in ^o^C (range) | | 37.1 (34.6-39.6) | 37.6 (34.3-39.3) | 0.1399 |
|  | Mean heart rate in bpm (range) | | 152.9 (111-190) | 175.4 (125-248) | **0.0069** |
|  | Mean respiratory rate in br/min (range) | | 45.9 (26-76) | 46.4 (28-70) | 0.8820 |
|  | Mean systolic blood pressure in mmHg (range) | | 89.9 (72-112) | 85.0 (63-120) | 0.2142 |
|  | Median saturation level in %O_2_ (range) | | 96.0 (42-100) | 96.0 (38-100) | 0.8402 |
| **Antibiotic treatment** | | |  |  |  |
|  | Oral antibiotic prescribed at discharge (Yes) | | 0 (0.0%) | 21 (84%) ^δ^ | **<0.0001** |
|  | Median duration of antibiotic treatment (range) | | 2.0 (2-3) | 10.5 (4-14) | **<0.0001** |
|  | Median number of antibiotics prescribed (range)^#^ | | 2 (2-3) | 3 (1-5) | **0.0005** |
| **Complications during antibiotic treatment** | | |  |  |  |
|  | Diarrhea (Yes) | | 8 (36.4%) | 11 (44.0%) | 0.3612 |
|  | Diaper rash (Yes) | | 10 (45.5%) | 12 (48.0%) | 0.5465 |
|  | Oral Thrush (Yes) | | 1 (4.5%) | 2 (8.0%) | 0.5498 |

**Abbreviations:** Kg, kilograms; h, hours; CTAS, Canadian Triage and Acuity Scale; ED, Emergency Department; bpm, beats per minute; br/min, breaths per minute; mmHg, millimetre of mercury; %O2, percentage of oxygen.

^Φ^ Text in bold denotes statistically significant differences (p<0.05) defined by unpaired t-test (numerical variables) or Chi-square (nominal variables).

^δ^ Four children from the long-term cohort were not prescribed any oral antibiotics at discharge, instead they received parenteral antibiotics for a total of 6 to 12 days of treatment.

^#^ Sum of antibiotic classes prescribed at ED and discharge.

Table S2: Permutation multivariance analysis of bacterial (shotgun) and fungal (ITS2) beta-diversity in ANTIBIO samples.

Data referent to Fig. S1e-f. Permutational multivariate analysis of variance stabilizing transformed community matrix testing the influence of study groups, time point, participant sex, age, pets, birth weight, and duration of antibiotic treatment over community matrix composition.

| **Whole Dataset** | | **Bacterial (Shotgun) Beta-Diversity Permutation** | | |  | **Fungal (ITS2) Beta-Diversity Permutation** | | |
| --- | --- | --- | --- | --- | --- | --- | --- | --- |
| **Variable** | **DF** | **F.Model** | **R^2^ (%)** | ***P*-value**^Φ^ |  | **F.Model** | **R^2^ (%)** | ***P*-value**^Φ^ |
| **Study group** | 1 | 1.4079 | 1.648 | 0.148 |  | 1.0104 | 1.232 | 0.425 |
| **Time point** | 1 | 4.4108 | 4.989 | <**0.001** |  | 1.0113 | 1.233 | 0.441 |
| **Patient sex** | 1 | 1.4035 | 1.643 | 0.158 |  | 1.0375 | 1.265 | 0.316 |
| **Patient age (days)** | 1 | 2.5865 | 2.987 | **0.008** |  | 0.8793 | 1.074 | 0.904 |
| **Pets at home** | 1 | 1.4943 | 1.748 | 0.124 |  | 1.0009 | 1.221 | 0.494 |
| **Birth weight** | 1 | 1.1772 | 1.382 | 0.305 |  | 1.1082 | 1.350 | 0.105 |
| **Days on antibiotic** | 1 | 1.5025 | 1.757 | 0.108 |  | 0.9706 | 1.184 | 0.563 |
| **Short-Term** |  |  |  |  |  |  |  |  |
| **Variable** | **DF** | **F.Model** | **R^2^ (%)** | ***P*-value**^Φ^ |  | **F.Model** | **R^2^ (%)** | ***P*-value**^Φ^ |
| **Time point** | 1 | 3.7265 | 2.849 | 0.373 |  | 0.9920 | 2.481 | 0.512 |
| **Patient sex** | 1 | 2.0646 | 5.424 | **0.021** |  | 1.1081 | 2.763 | 0.097 |
| **Patient age (days)** | 1 | 2.6043 | 6.746 | **0.009** |  | 1.0350 | 2.585 | 0.301 |
| **Pets at home** | 1 | 1.6099 | 4.28 | 0.079 |  | 1.0010 | 2.502 | 0.455 |
| **Birth weight** | 1 | 1.1246 | 3.029 | 0.320 |  | 1.1298 | 2.815 | 0.074 |
| **Days on antibiotic** | 1 | 2.1425 | 5.617 | **0.020** |  | 1.0001 | 2.500 | 0.467 |
| **Long-Term** |  |  |  |  |  |  |  |  |
| **Variable** | **DF** | **F.Model** | **R^2^ (%)** | ***P*-value**^Φ^ |  | **F.Model** | **R^2^ (%)** | ***P*-value**^Φ^ |
| **Time point** | 1 | 2.9391 | 6.006 | <**0.001** |  | 0.9827 | 2.398 | 0.565 |
| **Patient sex** | 1 | 0.5308 | 1.141 | 0.955 |  | 1.1763 | 2.857 | **0.045** |
| **Patient age (days)** | 1 | 0.6313 | 1.354 | 0.870 |  | 0.8635 | 2.113 | 0.948 |
| **Pets at home** | 1 | 1.4867 | 3.131 | 0.081 |  | 1.0352 | 2.523 | 0.332 |
| **Birth weight** | 1 | 2.2494 | 4.4662 | **0.002** |  | 1.2946 | 3.135 | **0.004** |
| **Days on antibiotic** | 1 | 1.2345 | 2.614 | 0.208 |  | 1.0934 | 2.661 | 0.140 |

^Φ^ Text in bold denotes statistical significance differences (*P*<0.05) defined PERMANOVA on Bray-Curtis dissimilarities.

Table S3: Differentially detected fungal ASV.

Data referent to Fig. S3e. Significant log2 FC detected in fungal ASV abundance in antibiotic-treated samples.

| **Short-Term** |  |  |  |  |
| --- | --- | --- | --- | --- |
| **ASV Number** | **Log2FC** | ***P-*value** | ***Padj*^Φ^** | **Species** |
| ASV_14 | 26.044334 | 5.00E-19 | 2.20E-17 | *s_Malassezia_globosa* |
| ASV_90 | 23.497566 | 9.15E-16 | 1.01E-14 | *s_Malassezia_restricta* |
| ASV_92 | 24.820430 | 2.02E-17 | 4.44E-16 | *s_Malassezia_sympodialis* |
| ASV_117 | 22.755141 | 7.18E-15 | 6.32E-14 | *s_Rigidoporus_sanguinolentus* |
| ASV_168 | 24.241520 | 1.10E-16 | 1.61E-15 | *s_Hyphodontia_pallidula* |
| **Long-Term** |  |  |  |  |
| **ASV Number** | **Log2FC** | ***P-*value** | ***Padj*^Φ^** | **Species** |
| ASV_2 | 8.6237383 | 2.87E-03 | 8.12E-03 | *s_Malassezia_sympodialis* |
| ASV_14 | 23.782171 | 4.37E-16 | 1.86E-15 | *s_Malassezia_globosa* |
| ASV_60 | 24.523258 | 5.16E-17 | 2.92E-16 | *s_Malassezia_restricta* |
| ASV_76 | 24.568051 | 4.52E-17 | 2.92E-16 | *s_Malassezia_globosa* |
| ASV_90 | 24.768774 | 2.51E-17 | 2.92E-16 | *s_Malassezia_restricta* |
| ASV_98 | 6.8703403 | 1.88E-02 | 3.99E-02 | *s_Fomitopsis_pinicola* |
| ASV_111 | 8.3356655 | 4.35E-03 | 1.06E-02 | *s_Resinicium_bicolor* |
| ASV_147 | 22.924533 | 4.82E-15 | 1.64E-14 | *s_Malassezia_restricta* |
| ASV_162 | 6.4945625 | 2.64E-02 | 4.98E-02 | *s_Rigidoporus_sanguinolentus* |

^Φ^ Significant differences defined by DESeq and FDR corrected; *Padj<*0.05.

Table S4: Determinants of *Malassezia* abundance trend identified by Linear Logistic Regression analysis.

Data referent to Fig. 1g. Logistic regression model of predictors of *Malassezia* abundance trend in ANTIBIO samples. Only the top five predictor factors identified by Random Forest analysis (data not shown) were tested for their association with the *Malassezia* abundance trend.

| **Factor** | **OR** | **95% CI** | ***P*-value**^Φ^ | **Code** |
| --- | --- | --- | --- | --- |
| Age_days | -0.0068 | (-0.0350 - 0.0212) | 0.621 | NS |
| Birth_weight | 0.6367 | (-0.9868 - 2.5729) | 0.472 | NS |
| Total_days_Abx | -0.1241 | (-0.4401 - 0.1159) | 0.356 | NS |
| Antibiotics_total_number | 2.1954 | (0.3604 - 4.9453) | **0.045** | * |
| Pregnancy_duration | -0.1941 | (-0.8197 - 0.3716) | 0.510 | NS |

**Abbreviations:** OR, log-transformed odds ratio; CI, 95% confidence interval.

^Φ^ Text in bold denotes statistically significant association (*P<*0.05).

Table S5: Network measures of centrality and hub taxa.

Data referent to Fig. S6. Jaccard’s similarity index (*j*) for measures of centrality (degree, betweenness, closeness, eigenvector) and hub taxa between sets of most central nodes between pre- and post-antibiotic networks. If *j=*0, the sets are completely different, and if *j=*1, the sets are exactly equal.

| **Measures of Centrality** | ***j*** | ***P(J ≤ j)***^Φ^ | **Code** |
| --- | --- | --- | --- |
| Degree | 0.220 | 0.0804 | ~ |
| Betweenness Centrality | 0.280 | 0.2612 | NS |
| Closeness Centrality | 0.208 | **0.0325** | * |
| Eigenvector Centrality | 0.208 | **0.0325** | * |
| Hub Taxa | 0.000 | **0.0034** | ** |

^Φ^ Text in bold denotes statistically significant *P(J ≤ j). P(J ≤ j)* determined by the probability of Jaccard’s index taking a value lower or equal to the calculated index *j.*

**Table S6: Volcano plot results for differentially detected metabolites.**

Data referent to Fig. 2f.

| **Compound** | **FC** | **Log2(FC)** | ***P*** ^Φ^ | **-log10(*P*)** |
| --- | --- | --- | --- | --- |
| Cortisol | 5.2990 | 2.4057 | 3.6589e-05 | 4.4367 |
| Adenine | 2.7548 | 1.4619 | 0.005525 | 2.2577 |
| Costicosterone | 2.2397 | 1.1633 | 0.007076 | 2.1502 |
| L-Ornithine | 0.4370 | -1.1941 | 0.009149 | 2.0386 |
| L-Arginine | 0.3424 | -1.5462 | 0.011048 | 1.9567 |
| N-Acetyl-L-aspartic acid | 4.5064 | 2.1720 | 0.022927 | 1.6396 |
| Oleate | 6.5824 | 2.7186 | 0.038268 | 1.4172 |

FC = fold change between B and B+M groups

^Φ^Statistically significant differences determined by unpaired T-test; *P<*0.05.
